# RAB14-dependent tubulovesicular recycling directs MET to invadopodia, promoting TNBC cell invasion

**DOI:** 10.1101/2025.09.23.677683

**Authors:** Amrita Khamari, Atreyee Guria, Kiran Tak, Shaily Verma, Rajiv Sharma, Yannis Kalaidzidis, Sunando Datta

## Abstract

Metastasis is one of the primary causes of cancer-related death in Triple-negative breast cancer patients. During metastatic dissemination, the cancer cells infiltrate in to surrounding tissue employing specialized membrane-protrusion with proteolytic activity called invadopodia. Cues from growth factors via the cognate receptor tyrosine kinases promote invadopodia formation and cancer cell invasion.

We report the role of HGF and its cognate receptor MET in TNBC invasion. The results from our study demonstrate that the TNBC cells when stimulated with HGF promotes invadopodia formation and facilitates delivery of MT1-MMP to the invadopodia in a MET dependent manner. We also showed that MET resides at invadopodia and its localization at invadopodia increases upon HGF treatment due to enhanced surface delivery of the RTK. By employing a degradation defective mutant, we demonstrated the critical role of MET recycling in driving breast cancer invasion. Moreover, we observed that the RCP–RAB14 axis mediates the surface delivery of MET through tubulovesicular carriers, with KIF16B promoting the formation of vesicular tubules on RAB14-positive endosomes.

Collectively, the study provides mechanistic insights on the growth factor-mediated recycling of RTK and the associated metalloprotease in invadopodia-dependent matrix degradation and cancer cell invasion.

## INTRODUCTION

Triple-negative breast cancer (TNBC) is a highly aggressive subtype of breast cancer and is characterized by the lack of expression of three key receptors: estrogen receptor (ER), progesterone receptor (PR), and human epidermal growth factor receptor 2 (HER2), which are generally targeted in conventional breast cancer therapies (1). The absence of these hormonal receptors in TNBCs imposes challenges in the treatment. In addition to the therapeutic challenges, TNBC exhibit a greater tendency for tumor invasion and metastasis which is one of the leading causes of higher mortality in TNBC patients (2,3). Metastasis requires the tumor cells to breach the tissue barrier employing proteolytic activity that remodel the extra cellular matrix (ECM), enabling the cancer cells to infiltrate in to the stroma (4). Subsequently the invasive cancer cells migrate to a distant organ through circulation and colonize to form a secondary tumor. Invadopodia, are actin-rich membrane protrusions decorated with proteases, that act as a tool for ECM and basement membrane degradation during cancer cell invasion (5). MT1-MMP is one of the invadopodia-associated metalloproteases and is overexpressed in TNBC cell lines. Perturbation in MT1-MMP delivery to invadopodia impairs ECM degradation and invasion (6,7). Studies suggest that the invasive cells in the leading front form invadopodia to create micro-tracks for the follower cells (8). The receptor tyrosine kinases (RTK), EGFR and MET are often amplified in TNBC patients (9) and can modulate the invadopodia-associated molecules, thereby promoting invadopodia-associated cancer invasion and metastasis (10–13).

Signaling cues from RTKs are essential for invadopodia formation and ECM remodelling (12,14). EGFR induces cortactin phosphorylation through SRC and Abl-related gene (ARG), which triggers the formation of invadopodia (10). However, MET-directed cortactin phosphorylation in invadopodia is SRC or ARG-independent, implicating for a distinct mode of function for MET in invadopodia formation (12). A chemically induced cytoplasmic variant of the RTK, TPR-MET disrupts the balance between intracellular signalling and regulatory mechanisms, promotes invadopodia associated ECM degradation (12,15,16).

At the cell surface, the RTKs bind to their respective ligands, leading to receptor dimerization and activation. Subsequently, they internalize rapidly and continue to signal from endosomes (17). The signalling of RTKs from the endosomes are tightly regulated by endocytic trafficking (18). From the early endosomes, the RTKS are sorted in to distinct pathways directing them either towards lysosomal degradation or recycling to deliver them back to the plasma membrane (17,19). The sorting of cargo from the early endosomes is a highly co-ordinated process, that is majorly facilitated by RAB GTPases (20,21). RAB GTPases act as molecular switches, that provide membrane identity by recruitment of downstream effectors. RABs recruit tethering complexes, molecular motors, adapters for cargo or motors, lipid modifying enzymes etc, together which promotes efficient cargo selection, segregation and sorting (22). Membrane tubulation is a key mechanism of cargo sorting on the endosomal membrane (23,24). The tubular membrane curvature are often initialled and stabilized by Sorting nexins (SNX) having a BAR domain (25). The pulling force exerted by the molecular motors further help to elongate the structure (26–28). RAB dependent recruitment of specific machinery ensure the enrichment of cargo at these microdomains. Upon fission these tubular structures becomes independent carrier for cargo trafficking (29).

RAB GTPases like RAB4, RAB11 along with their associated trafficking machinery are known to facilitate the recycling of MET to the plasma membrane driving cell migration (30,31). Additionally, SNARE-dependent delivery of EGFR to invadopodia promotes invadopodia-associated tumor invasion, highlighting the potential role of RTK trafficking in cancer cell invasion (11). Although cell migration and invasion are tightly coupled processes, the significance of MET trafficking in cancer cell invasion is still poorly understood.

Here, we investigated the role of HGF and its cognate receptor, MET, in invadopodia-associated cancer cell invasion in TNBC cell lines. Our study revealed that activated MET resides at invadopodia and its invadopodia-recruitment is enhanced upon HGF stimulation. Our findings emphasize the importance of the growth factor-mediated cell-surface transport of MET in invadopodia formation and MT1-MMP recruitment through physical interaction with the protease that leads to enhanced TNBC invasion upon HGF stimulation. We further delineated the involvement of the cellular recycling repertoire in the growth factor-stimulated recycling of MET.

## RESULTS

### HGF-induced MET activity promotes TNBC breast cancer invasion through invadopodia-mediated ECM degradation

The upregulation of EGFR and MET promotes breast cancer invasion (32,33). We examined the MET expression in TNBC cell lines MDA-MB-231, BT-549, and the pre-invasive MCF10A DCIS by immunoblotting. We observed comparable MET expression in all 3 cell lines (Fig. S1A, S6). MET-HGF signalling axis is reported to promote invasion in gastric cancer cells and melanoma cells (13,34). To inspect the effect of HGF on the invasive properties of breast cancer cell lines, we employed a spheroid invasion assay that closely resembles the physiological environment. Spheroids were formed using MDA-MB-231 or BT-549 cells and embedded in collagen I with or without HGF or in complete media. The spheroids were allowed to invade for 48 hours. The ratio of core area to invaded area was calculated, revealing that HGF-treated cells invaded more into collagen compared to untreated conditions (Fig. 1A, S1B, MOVIE 1).

**Figure 1:**
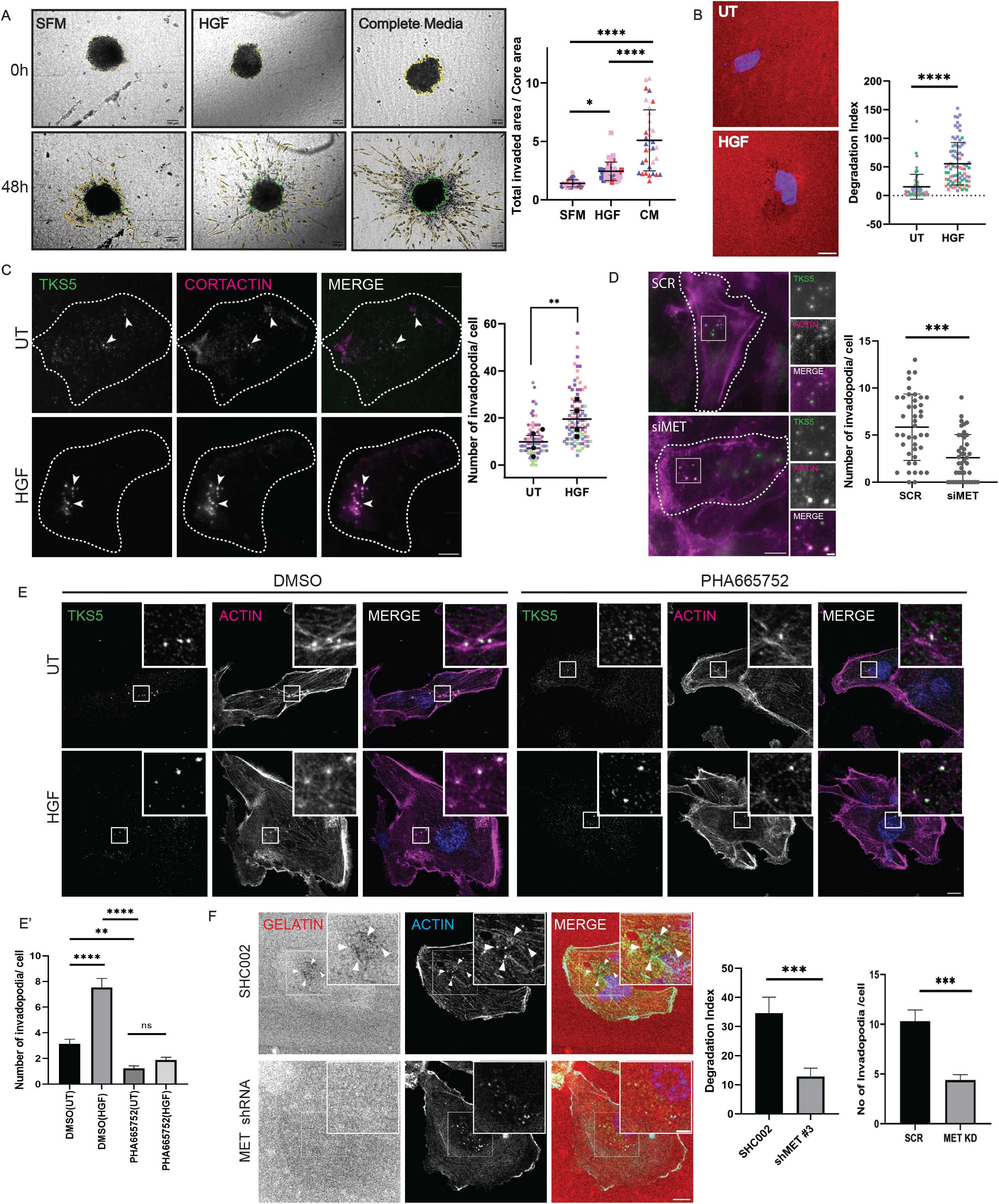
HGF-induced MET activity promotes TNBC breast cancer invasion through invadopodia-mediated ECM degradation. (A) Bright-field images of MDA-MB-231 spheroids embedded in the collagen matrix at 0h and 48h in the presence or absence of HGF or complete media. The ratio of the invaded area to the core area was quantified and plotted. The core has been outlined in green, and the invaded area has been outlined in yellow. Data points from individual biological replicates are distinctly color-coded. The error bar represents Mean ±SD, One-way ANOVA with multiple comparisons. **** P < 0.0001, * P < 0.05, Scale bar: 100µm. N=3, n=30. (B) MDA-MB-231 cells were seeded on Alexa568-labeled gelatin-coated coverslip with or without HGF for 3h. Cells were fixed and imaged with a confocal microscope. The degradation index was quantified and plotted. The arrowhead represents degradation spots. Data points from individual biological replicates are distinctly color-coded. The error bar represents Mean ±SD, Unpaired t-test, **** P < 0.0001. scale bar: 10µm. N=3, n=200. (C) MDA-MB-231 cells were seeded on a gelatin-coated glass-bottom dish and allowed to attach for 3 hours, followed by incubation with or without HGF for an additional 3 hours. Cells were fixed and immunostained for TKS5 and cortactin. Images were captured using a TIRF microscope. TKS5 puncta positive for cortactin were considered as invadopodia. Arrowheads indicate TKS5-cortactin positive puncta. Data points from individual biological replicates are distinctly color-coded. The error bar represents Mean ±SD, Paired t-test, ** P < 0.01. scale bar: 10µm. N = 4, n = 150. (D) MDA-MB-231 cells were transfected with control or MET siRNA. After 60 hours of transfection, cells were seeded on a gelatin-coated glass-bottom dishes for 6h. Cells were fixed and immunostained for TKS5, and F-actin. Images were captured using a Nikon TIRF microscope. TKS5 and Actin-positive structures were identified and plotted. The inset shows a magnified view of the region indicated by the box. The error bar represents Mean ±SEM, Unpaired t-test, *** P < 0.001. scale bar: 10µm. Inset: 2µm. N=3, n=60. (E, E’) MDA-MB-231 cells were seeded on gelatin-coated coverslips for 3h, followed by incubation with vehicle or PHA665752 (5 µM) with or without HGF for an additional 3h. Cells were fixed, immunostained with anti-TKS5 antibody, Phalloidin to stain F-actin, and DAPI. Images were captured using a confocal microscope. TKS5 puncta positive for Actin were considered as invadopodia. The inset shows a magnified view of the region indicated by the box. scale bar: 10µm. Inset: 2µm. The error bar represents Mean ±SEM, One-way ANOVA with multiple comparisons. **** P < 0.0001, ** P < 0.01, ns P > 0.05, N=3, n=200. (F) MDA-MB-231 cells were transfected with control SHC002 or MET shRNA clone #3, and after 36h of transfection, cells were seeded on Alexa568-labeled gelatin-coated coverslip for 6h. The cells were fixed and immunostained with anti-MET antibody, Phalloidin, and DAPI. Degradation index and degradation spots immunostained for Actin were quantified. The inset shows a magnified view of the region indicated by the box. The Arrowheads indicate the degradation spots that were immunostained for Actin. scale bar: 10µm. Inset: 2µm. The error bar represents Mean ±SEM, Unpaired t-test. *** P < 0.001, N=3, n=200.

To invade the stroma, tumor cells degrade the underlying basement membrane. To understand the effect of HGF on ECM remodelling, we carried out a fluorescent-labelled gelatin degradation assay. We seeded cells on Alexa568-labeled gelatin-coated coverslips in the presence or absence of HGF and allowed them to degrade gelatin for a minimum of three hours. The loss of fluorescence due to gelatin degradation, represented as the degradation index, was calculated as detailed in the methods section. Our results showed that HGF treatment significantly increased gelatin degradation activity in MDA-MB-231, BT-549, and MCF10A DCIS cell lines (Fig. 1B, S1D-E).

The ECM degradation activity of cancer cells is attributed to invadopodia formation (35,36). Therefore, we investigated the ability of HGF to induce invadopodia in these cell lines. Cells were seeded on gelatin in the presence or absence of HGF for 6h, fixed, and immunostained for known invadopodia markers, i.e., TKS5, cortactin, or Actin. Puncta positive for two invadopodia markers were considered as invadopodia. When stimulated with HGF, increased numbers of invadopodia were observed in MDA-MB-231, BT-549 and MCF10A DCIS cells (Fig. 1C, S1D, F). The percentage of cells forming invadopodia was also found to be increased upon HGF treatment in MDA-MB-231 cells (Fig. S1G). Additionally, the other most abundant RTK in TNBC cells, EGFR, when activated by EGF, resulted in a significant increase in invadopodia number in MDA-MB-231 (Fig. S1H) cells, corroborating previous studies (10).

To investigate, whether MET is important for HGF-governed invadopodia formation, we depleted MET using siRNA and TKS5-Actin positive structures were quantified. The invadopodia numbers were found to be decreased upon MET silencing (Fig. 1D, S1I, S6). Further, to substantiate this observation, we used 2 individual siRNA targeting MET (Oligo#1, #2). Consistent with the SMARTpool results, MET silencing using individual oligoes resulted in an approximate 50% reduction in the invadopodia numbers (Fig. S1C). The percentage reduction in gelatin degradation in cells treated with SMARTpool, Oligo#1 or Oligo#2 were also comparable (Fig. S1C). Since MET is an RTK, we asked whether the activation of MET is essential for invadopodia formation. To address this question, we inhibited the kinase activity of MET using the drug PHA665752. Immunoblot analysis revealed that HGF stimulation increased MET pTyr1234-1235 (p-MET), which was abolished by the drug (Fig. S1J). Interestingly, the drug treatment also promoted degradation of MET (Fig. S1J, S6). MDA-MB-231 cells were analyzed for their ability to form invadopodia in the presence of 5 µM of the drug or vehicle, with or without HGF. Inhibition of MET kinase activity using PHA665752 abolished the HGF-induced invadopodia formation (Fig. 1E, E’).

Next, we silenced MET using shRNA and analyzed the the effect on gelatin degradation. Different clones of shRNA against MET were transfected, and the level of silencing was evaluated (Fig. S1K, S6). MDA-MB-231 cells were transfected with control SHC002 or MET shRNA C#3. After 36h, cells were seeded on labelled gelatin for 6h, fixed, and immunostained with phalloidin and an antibody against MET. Gelatin degradation and invadopodia formation was abrogated in the MET-depleted cells compared to the control (Fig. 1F, S1L).

Taken together, HGF activates RTK MET and promotes TNBC invasion by enhancing invadopodia formation and associated activity.

### Enhanced MET recruitment to invadopodia upon HGF stimulation

Since MET activity was found to be important for invadopodia formation, we first asked if MET is localized at invadopodia. MDA-MB-231 cells were seeded on a gelatin-coated surface for 6 hours. Then, the cells were fixed and immunostained for MET and cortactin. A notable population of cortactin puncta was found to be localized with MET (Fig. 2A). Next, the colocalization of MET to GFP-TKS5 was examined using TIRF microscopy (TIRFM). Approximately 30% of the TKS5 population showed staining for MET (Fig. 2B, D). Confocal imaging of BT-549 cells immunostained for TKS5 and MET revealed that MET-TKS5 colocalization follows the same trend as MDA-MB-231 (Fig. S2A). Next, we asked whether MET is in its active form at invadopodia. Cherry-TKS5 expressing cells were seeded on gelatin, fixed, and immunostained for phosphorylated MET. TIRF imaging revealed that 27% of TKS5 puncta are positive for p-MET, similar to that of MET localization to TKS5-positive invadopodia (Fig. 2C, D).

**Figure 2:**
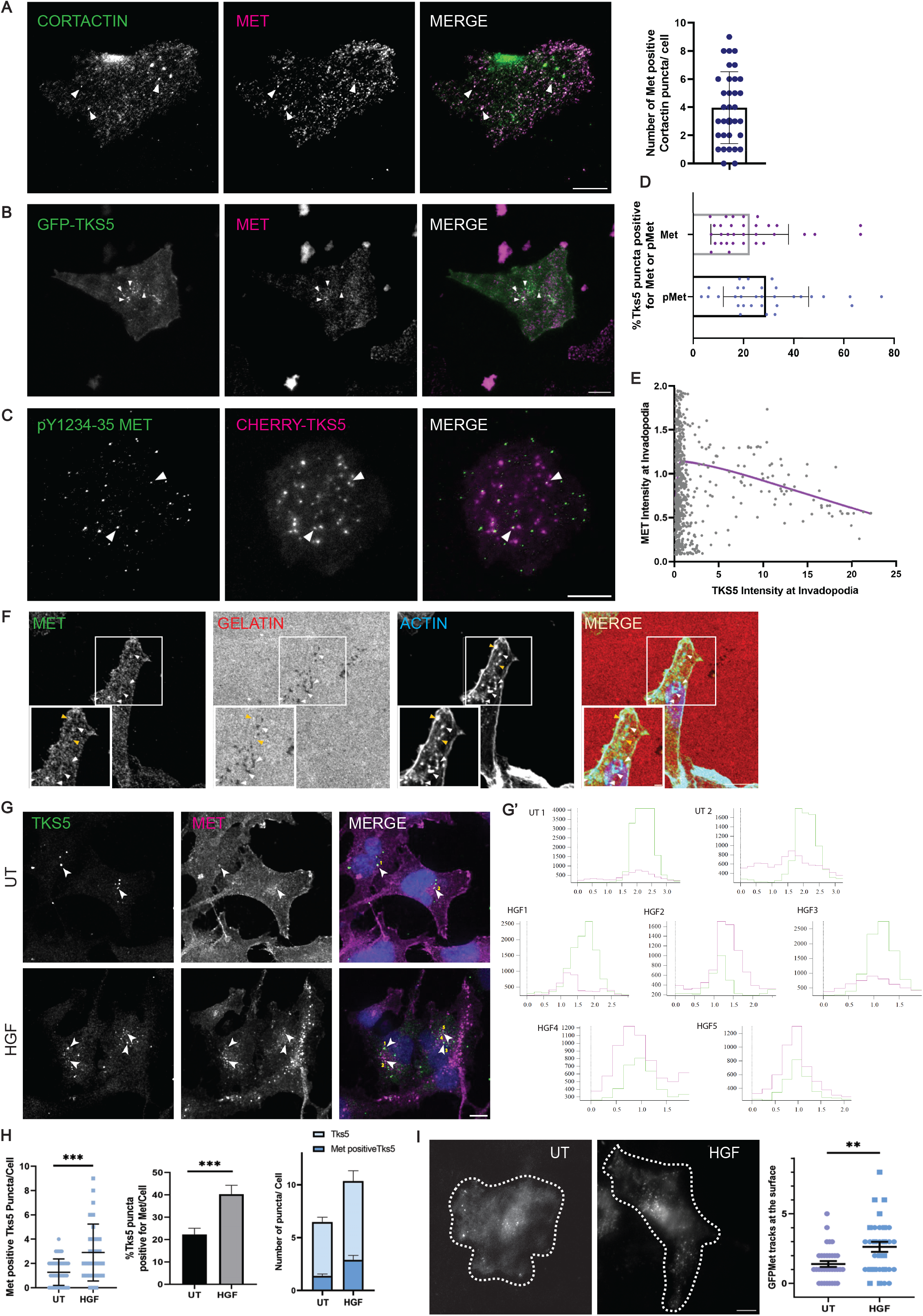
Enhanced MET recruitment to invadopodia upon HGF stimulation. (A) MDA-MB-231 cells were seeded on gelatin-coated imaging dishes for 6h. Cells were fixed and immunostained for cortactin and MET. Images were captured using a TIRF microscope. Colocalized cortactin-MET puncta were quantified and plotted. The arrowheads indicate the MET-positive cortactin puncta. The error bar represents Mean ±SD, scale bar: 10µm, n= 60. (B) GFP-TKS5-expressing MDA-MB-231 cells were seeded on gelatin-coated imaging dishes for 6h. Cells were fixed, immunostained for MET and imaged with TIRFM. The arrowhead indicates the MET-positive TKS5 puncta. scale bar: 10µm. (C) Cherry-TKS5-expressing MDA-MB-231 cells were seeded on gelatin-coated glass-bottom imaging dishes for 6h. Cells were fixed, immunostained with anti-p-MET antibody and imaged with TIRFM. The arrowhead indicates the p-MET-positive TKS5 puncta. scale bar: 10µm. (D) Colocalized TKS5-MET or p-MET puncta were quantified and plotted. The error bar represents Mean ±SD, N=3, n ≤ 50. (E) The fluorescence intensity of TKS5 and MET at individual invadopodium was quantified using Motiontracking and normalized with their respective mean intensities. The normalized fluorescence intensities of TKS5 (X-axis) and MET (Y-axis) were plotted. The points were fitted with a smoothing spline. n=800 (invadopodia). (F) MDA-MB-231 cells were seeded on Alexa568-labeled gelatin-coated coverslips for 12h. Cells were fixed and immunostained for F-actin and MET. The inset shows a magnified view of the region indicated by the box. The white Arrowhead represents Actin at the degradation spots, and the yellow arrowhead represents Actin-MET puncta without degradation. scale bar: 10µm. Inset: 2µm. (G) BT-549 cells were seeded on gelatin-coated coverslips for 3h followed by incubation with or without HGF for 3h. Cells were fixed, immunostained for TKS5, MET and Nucleus. Images were taken using the confocal microscope. scale bar: 10µm. (G’) Using Motiontracking a line was drawn across indicated MET-TKS5 puncta and the intensities of both channels were plotted. (H) Colocalized TKS5-MET puncta were quantified using Motiontracking and plotted. The error bar represents Mean ±SEM/SD, Unpaired t-test, *** P < 0.001, N=3, n = 200. (I) BT-549 cells expressing GFP-MET were seeded on glass-bottom dishes. The cells were imaged using a TIRF microscope in the presence or absence of HGF. The GFP-MET vesicles moving in the TIRF field for a minimum of 4 consecutive frames were quantified and plotted as MET tracks. The error bar represents Mean ±SEM, Unpaired t-test. scale bar: 10µm. ** P < 0.01, N=3, n=30.

TKS5 is a critical component of invadopodia and its concentration gradually increases as the invadopodia matures (37). When we plotted the normalized steady-state fluorescence intensity of MET and TKS5 localized at invadopodia, the resulting plot showed two different populations of invadopodia. Invadopodia with TKS5 intensity less than 4 showed no correlation with MET intensity (R^2^ = 0.03 for linear regression); however, invadopodia with TKS5 intensity more than 4 showed a negative correlation (R^2^ = 0.3 for linear regression) (Fig. 2E). Further, we analyzed the frequency distribution of the MET to TKS5 fluorescence intensity ratio across invadopodia. It revealed that most invadopodia structures exhibited lower MET/TKS5 ratios (Fig. S2B). This observation prompted us to investigate the apparent absence of MET from mature invadopodia. To address this, the localization of MET at the invadopodia-mediated degradation spots was examined in the MDA-MB-231 cells plated on fluorescence-labelled gelatin. As implied above, the degradation spots did not show strong colocalization with MET (Fig. 2F).

As we have observed that HGF stimulates invadopodia formation via MET and the RTK is physically localized at the invadopodia site, we asked if HGF influences MET localization to the invadopodia. Cells were seeded on gelatin with or without HGF, fixed, and immunostained for MET and invadopodia markers. Intriguingly, upon treatment with HGF, we observed an increased MET-containing TKS5-positive invadopodia structure in both MDA-MB-231 and BT-549 (Fig. 2G-H, S2C-D). This observation led us to hypothesize that HGF may induce the surface delivery of MET to invadopodia.

Previously, it was reported that HGF stimulation leads to endocytosis of MET, followed by lysosomal degradation of the RTK (16,38). First, we analyzed the effect of HGF on receptor endocytosis and degradation of MET. To study the HGF-mediated receptor endocytosis, the colocalization of MET with RAB5 was quantified with or without HGF. The percent colocalization of MET with RAB5 was found to be increased from 12% to 43% in the presence of HGF stimulation (Fig. S2E). Next, cells were incubated with HGF for different time points, and the total MET levels were quantified using immunoblotting. After 3h of HGF stimulation, ∼44% of MET was found to be degraded (Fig. S2F, S6).

Next, to validate our hypothesis that HGF can induce surface delivery of MET, GFP-MET-expressing cells were imaged with TIRFM with or without HGF, and MET tracks were quantified as explained in the method. In the presence of HGF, an increased number of MET tracks were found near the plasma membrane compared to the untreated cells (Fig. 2I, MOVIE 2). This suggests that HGF indeed increases the trafficking of MET to the cell surface. We also observed a similar effect of EGF on EGFR localization at invadopodia (Fig. S2G, H), consistent with an earlier study in MDA-MB-231 (11).

These observations convey that MET resides in the invadopodia structures and HGF treatment leads to enhanced surface delivery of MET and its recruitment to invadopodia.

### HGF-mediated surface delivery of MET promotes invadopodia-associated ECM degradation and breast cancer cell invasion

As HGF promotes MET recycling and breast cancer invasion, we investigated whether MET recycling drives invadopodia-associated TNBC invasion. To address this, we employed the degradation-defective M1250T mutant and the endocytosis-defective LL/AA MET mutant as a negative control. An oncogenic mutation in MET at M1268T (M1250 in the MET constructs used in this study) is naturally found in papillary renal carcinoma, has a degradation defect leading to continuous endocytosis and recycling (31). The dileucine motif of MET is required for endocytosis of the receptor, which is mutated to AA in this study (39). First, we have characterized the mutants for endocytosis and degradation. We looked into the degradation kinetics of the mutants by immunoblotting. Unlike the wild-type (WT) MET, the M1250T MET mutant did not undergo degradation upon HGF stimulation (Fig. 3B, S6). Since the M1250T mutant has a slower degradation rate, we asked whether the M1250T mutant is in its activated form. V5 tagged-MET WT or the mutant was overexpressed and pulled down using Ni-NTA beads through the His-tag present in the coding region. The pulled-down samples were analysed by immunoblotting using the p-MET antibody. M1250T MET was highly phosphorylated at the active site compared to the WT MET (Fig. 3C, S6). The observation was further confirmed by immunofluorescence. The normalized CTCF value of p-MET in M120T MET expressing cells was found to be higher compared to WT MET (Fig. S2I). Next, the effect of HGF on the endocytosis of WT or the mutant MET was investigated. Cells were transfected with WT V5-MET or the mutant constructs, and their percentage of localization with EEA1, an early endosomal marker, was quantified in the presence or absence of HGF. HGF stimulation leads to an increased MET population localizing with EEA1 in WT MET expressing cells, whereas M1250T mutant cells showed an increased colocalization with EEA1 irrespective of the HGF stimulation (Fig. 3D, D’). The endocytosis-defective LL/AA mutant did not show any increase in the EEA1 colocalization in response to HGF.

**Figure 3:**
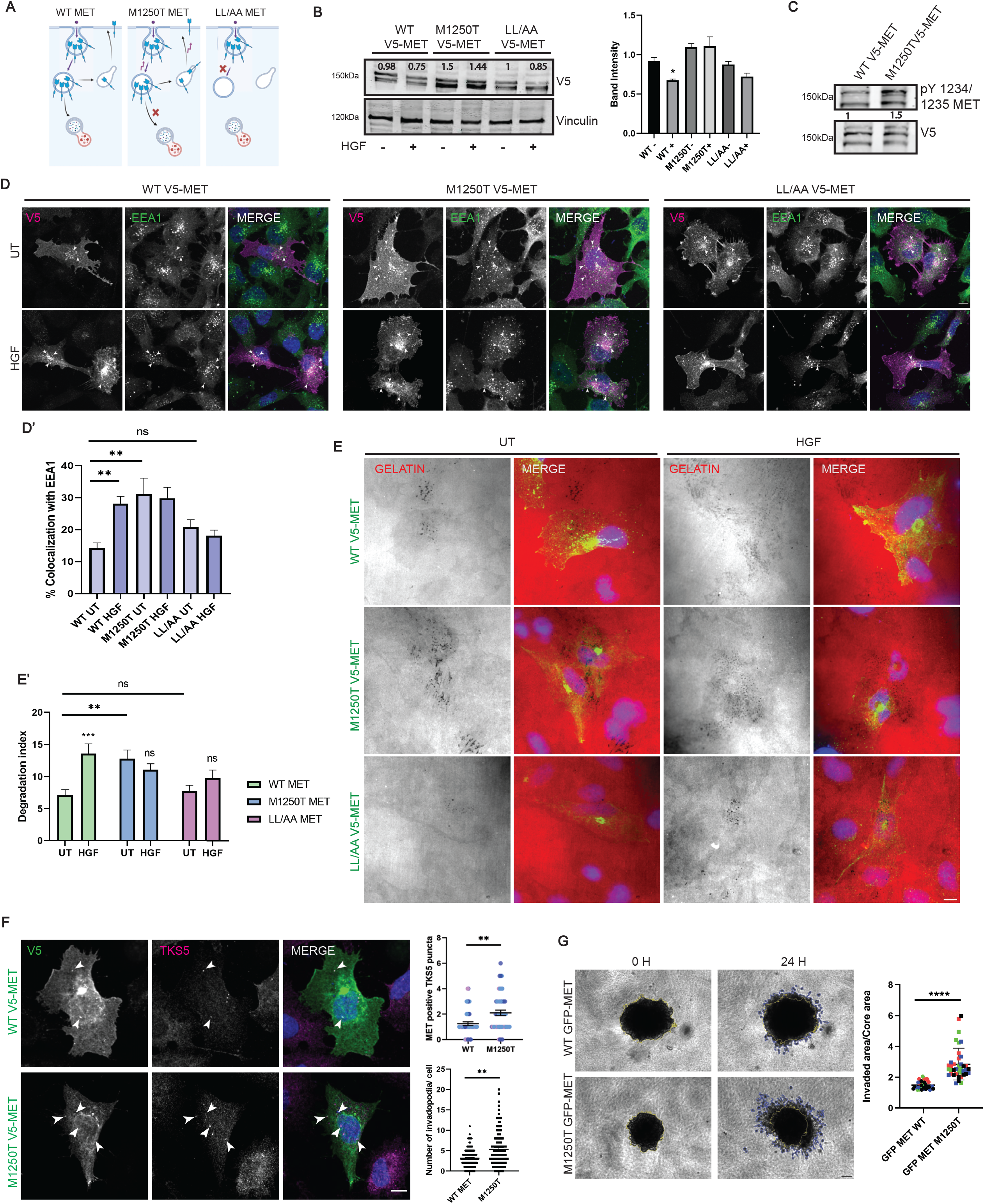
Intracellular trafficking of MET promotes invadopodia-associated ECM degradation and breast cancer cell invasion. (A) Graphical representation of WT MET and mutants’ endocytosis and degradation. (B) BT-549 cells expressing WT or the mutants of V5-MET were treated with or without HGF for 2 hours, followed by cell lysis. The lysates were subjected to immunoblotting with anti-V5 and anti-vinculin antibodies. The V5 signal intensity was quantified and normalized with Vinculin. Error bar represents Mean ±SEM. Unpaired t-test. * P < 0.05. (C) BT-549 cells expressing WT or M1250T V5-MET were lysed and incubated with Ni-NTA beads. The pulled-down samples were separated by SDS PAGE and processed for immunoblotting with anti-V5 and anti-p-MET antibody. (D, D’) WT or mutant V5-MET expressing BT-549 cells were seeded on glass coverslips with or without HGF. Cells were fixed, immunostained for V5 and EEA1. Images were captured with a confocal microscope. V5-MET puncta localizing with EEA1 were quantified and plotted. scale bar: 10µm. The error bar represents Mean ±SEM. ** P < 0.01. ns P > 0.05 N=3, n=60. (E, E’) BT-549 cells expressing V5-MET WT or mutants were seeded on Alexa568-labeled gelatin-coated coverslips with or without HGF for 6h. Cells were fixed, then immunostained for V5 and the Nucleus and imaged with the confocal microscope. Gelatin degradation by the V5-MET overexpressing cells was calculated and plotted. scale bar: 10µm. The error bar represents Mean ±SEM, Unpaired t-test. ** P < 0.01, ns P > 0.05, N=3, n=120. (F) V5-MET WT or M1250T mutant expressing BT-549 cells were seeded on a gelatin-coated coverslip for 6h and fixed. Cells were immunostained for V5, TKS5, and images were captured with the confocal microscope. The error bar represents Mean ±SEM, Unpaired t-test. scale bar: 10µm. ** P < 0.01, N=3, n=120 (G) Bright-field images of GFP-MET WT or M1250T expressing BT-549 spheroids embedded in the collagen matrix at 0h and 24h. The ratio of the invaded area to the core area was quantified and plotted. Data points from individual biological replicates are distinctly color-coded. The error bar represents Mean ±SD, Unpaired t-test. scale bar: 100µm. **** P < 0.0001, N=3, n=40.

To understand the effect of intracellular trafficking of MET on breast cancer invasion, cells expressing the WT or mutant forms of MET were analyzed for their ECM degradation ability. WT or the mutant V5-MET expressing BT-549 cells were seeded on Alexa568-labeled gelatin-coated coverslip with or without HGF, fixed, and immunostained with anti-V5 antibody. Confocal image analysis showed that WT MET-expressing cells have enhanced ECM degradation ability when stimulated with HGF; however, neither mutant responded to HGF (Fig. 3E, E’). The mutant M1250T MET showed comparable gelatin degradation as the WT MET stimulated with HGF, whereas the LL/AA had decreased invasion ability. We repeated the experiment in MDA-MB-231 cells and observed similar trends for the mutants (Fig. S2J).

Since the ECM degradation ability of M1250T was higher, we asked whether it was because of increased invadopodia formation. M1250T MET-expressing cells showed increased invadopodia formation compared to WT MET-expressing cells (Fig. 3F, S2K, K’). However, cells expressing the LL/AA mutant did not show a significant difference compared to WT MET (Fig. S2K, K’). Also, the number of invadopodia positive for M1250T MET was higher compared to WT MET-positive invadopodia (Fig. 3F)

To further investigate the effect of the M1250T MET mutant on breast cancer invasion, we employed a spheroid invasion assay. We generated a doxycycline-inducible GFP-MET WT and M1250T cell line in BT-549. Spheroids were formed for WT and M1250T MET expressing cells and embedded in collagen for 24 hours to allow invasion. The M1250T GFP-MET expressing cell line invaded more into the collagen compared to the WT MET expressing spheroid (Fig. 3G), supporting our observations from gelatin degradation.

These observations demonstrated that MET recycling ensures a steady delivery of the RTK to the invadopodia structures, thereby promoting breast cancer invasion.

### HGF promotes RAB4 & RAB14-mediated MET delivery to the plasma membrane

RAB GTPases are molecular switches, localized to distinct endosomal compartments, and orchestrate various endocytic, trans, and exocytic pathways (40). To investigate the RABs involved in MET recycling, we first inspected the colocalization of MET with endosomal markers that work in the recycling pathways, like RAB4, RAB22, RAB14 and RAB11. Cells were transfected with the different RAB constructs and incubated in the presence or in the absence of HGF before fixing and immunostaining for MET. The object-based percentage of colocalization was calculated using Motiontracking, for which vesicles for each molecule were identified, and the objects were considered to be colocalized if the overlap of their respective area was more than 35%. While the localization of MET with RAB4 and RAB14 was found to be increased in the HGF-stimulated cells (Fig. 4A-C, S3A-C), the colocalization with RAB22 and RAB11 remained unchanged, suggesting them to be associated with an HGF-independent trafficking pathway (S3D, E).

**Figure 4:**
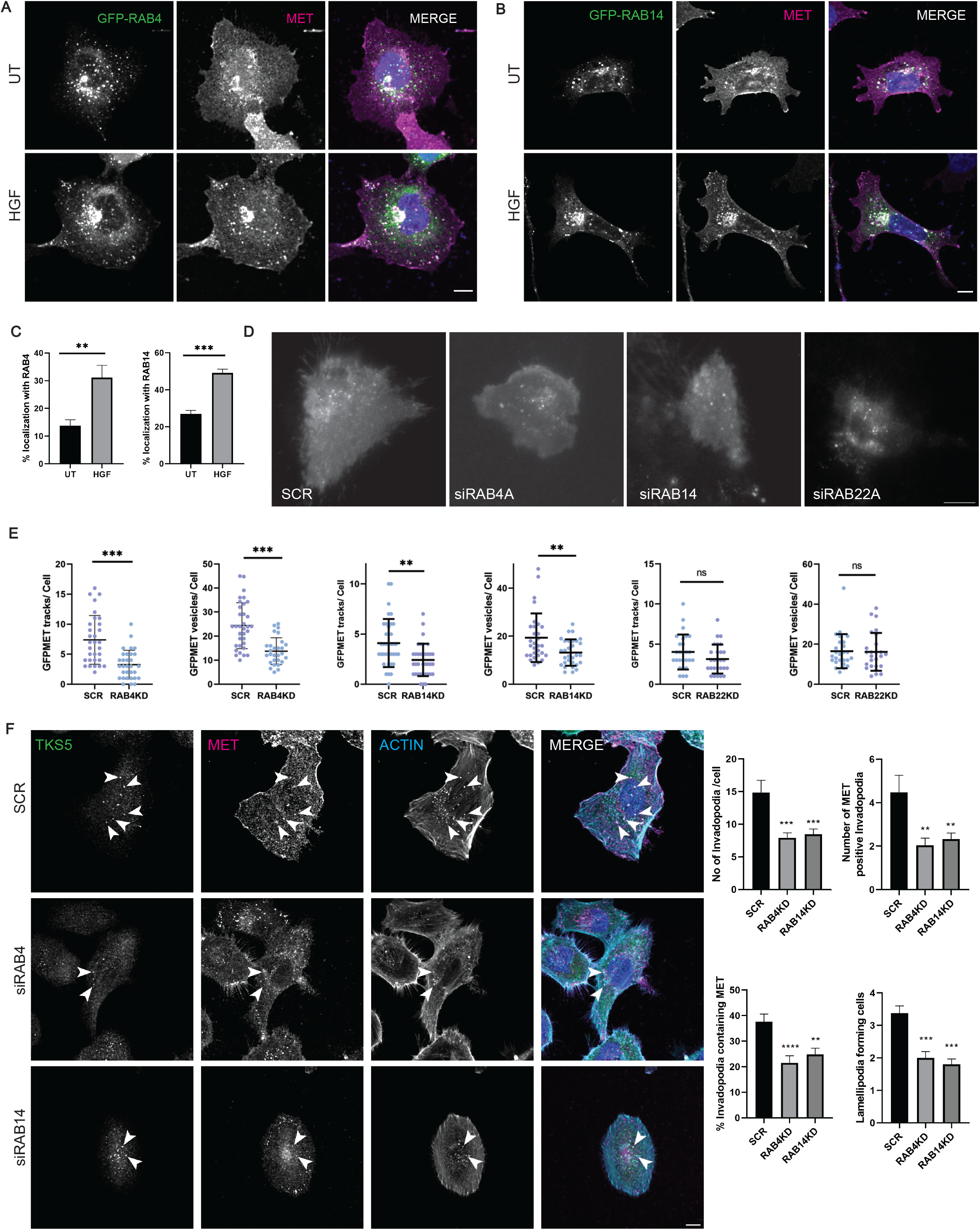
HGF promotes RAB4 & RAB14 mediated MET delivery to the plasma membrane. BT549 cells, seeded on glass coverslips, were transfected with (A) GFP-RAB4 or (B) GFP-RAB14. After 14h of transfection, cells were incubated for 2h with or without HGF, then fixed and immunostained for MET. Images were captured using a confocal microscope. scale bar: 10µm. (C) Graph showing the percent colocalization of MET with GFP-RAB4 and GFP-RAB14 in the presence of absence of HGF treatment. The error bar represents Mean ±SD, Unpaired t-test. ** P< 0.01. N=3, n=90 (D) BT-549 cells were transfected with siRNA against RAB4, RAB14, RAB22, or control siRNA. After 60 hours, cells were transfected with GFP-MET and seeded on the gelatin-coated glass-bottom dish. The GFP-MET-transfected cells were imaged in the presence of HGF using a TIRF microscope for 30 frames without an interval. scale bar: 10µm (E) The GFP-MET vesicles forming the track were identified as mentioned in the Method section. The numbers of GFP-MET tracks moving for at least 4 consecutive frames were quantified for each condition and plotted. In addition, the total number of GFP-MET vesicles was quantified. The error bars represent Mean±SD. Unpaired t-test. *** P< 0.001, ** P< 0.01, ns P > 0.05, N=3, n = 30. (F) BT-549 cells were depleted with RAB4 or RAB14, and after 60h, cells were seeded on gelatin-coated coverslips for 6h with 3h of HGF stimulation. Cells were fixed and immunostained for MET, TKS5, and F-actin. Images were captured in a confocal microscope. MET colocalizing with invadopodia was quantified and plotted. The number of cells forming lamellipodia was quantified. scale bar: 10µm. The error bar represents Mean ±SEM, One-way ANOVA, *** P< 0.001, ** P< 0.01. N=3, n=200.

Next, to identify the RABs that are involved in the HGF-dependent MET surface delivery, live cell TIRF imaging was performed in RAB4, RAB14, and RAB22-depleted cells. As the role of RAB4 in MET recycling has been established before, it was included as a positive control (30). The RABs were silenced using SMARTpool siRNA, and knockdown efficiency was quantified (Fig. S3F). GFP-MET was overexpressed in the control and the RABs-depleted cells and imaged using a TIRF microscope in the presence of HGF. The total number of MET vesicles, including MET tracks at the surface, was found to be reduced when RAB4 or RAB14 was deleted compared to the control cells (Fig. 4D-E, MOVIE 3).

When the recycling is impaired, a significant population of the RTKs is subjected to degradation (31,41). Next, to study the fate of MET in the RABs-depleted cells, we performed immunoblotting to quantify the total population of cellular MET. Cells depleted of the RABs mentioned above were treated with HGF. Cells were lysed and subjected to immunoblotting. The RAB4 and RAB14-depleted cells were observed to have a decreased total MET level compared to the control (Fig. S3G, S6).

Since the retromer is known to drive the recycling of various cargoes to the plasma membrane, we checked the role of retromer depletion on MET recycling (42). The co-localization of MET with VPS35, a retromer subunit, was comparable with or without HGF (Fig. S3M). Further to analyze whether retromer is essential for MET trafficking, MDA-MB-231 cells were depleted for the VPS26A retromer subunit, and the MET localization to Actin at fan-shaped membrane protrusions in response to HGF stimulation was quantified. The membrane pool of MET localizing with actin-rich structures was unchanged in the retromer-depleted condition (Fig. S3N). The total MET in the retromer-silenced cells also remained unaffected (Fig. S3K).

As HGF stimulation leads to increased recruitment of MET to invadopodia, we asked whether RAB4 & RAB14-mediated MET recycling plays a role in invadopodia formation and MET colocalization at the invadopodia. Cells depleted with RAB4 or RAB14 were analyzed for their ability to form invadopodia and the MET localization to the structure. As expected, invadopodia numbers were reduced in RAB4 and RAB14-depleted cells. Further, MET localization to invadopodia was also found to be reduced (Fig. 4F). In line with previous findings, the number of lamellipodia-forming cells was also drastically reduced in the RAB4 and RAB14-silenced cells (Fig. 4F) (30,41,43).

We examined the colocalization of RAB14 with EEA1, and 60% of the RAB14 was found to be colocalized with EEA1 (Fig. S3H). However, a small population of MET-containing RAB14 vesicles (∼10%) was not EEA1 positive and present away from the nucleus. Next, we analyze the distribution of MET to RAB4 and RAB14 vesicles using super-resolution microscopy. The GFP-RAB4 and Cherry-RAB14 expressing cells were stimulated with HGF, then fixed and immunostained for MET. The peripheral vesicles containing MET were positive for RAB4 or RAB14, with RAB14-positive MET endosomes showing minimal or no RAB4 presence, whereas the RAB4-positive MET endosomes are devoid of RAB14 (Fig. S3I).

A study from Wang et.al. suggests that RAB14 vesicles can also be budded out from the Golgi (44). We analyzed the localization with a Golgi marker to understand the biogenesis of these RAB14-MET vesicles. Cherry-RAB14 expressing cells were fixed and immunostained with anti-MET and anti-Golgin97 antibody. A small fraction (4%) of RAB14-positive MET puncta localized with the Golgi (Fig. S3L).

When we analyzed the effect of HGF on RAB14 vesicular characteristics. GFP-RAB14 expressing cells were treated with or without HGF, fixed, and imaged with a confocal microscope. The integral intensity and area of RAB14 vesicles were quantified. The area of GFP-RAB14 vesicles was found to be increased in the HGF-treated cells, whereas the integral intensity was unaffected (Fig. S3J). This indicates that the effect of HGF on the trafficking pathway is not mediated solely by RAB14, but also that additional players are involved in the RAB14-mediated trafficking.

The above results suggest that a fraction of the MET recycles back to the surface via the RAB4 or RAB14-mediated recycling pathway, which is crucial for the recruitment of MET to invadopodia.

### RCP-RAB14-KIF16B axis regulates MET trafficking

The mechanism behind HGF-mediated MET recycling through RAB4 has been demonstrated earlier (30,41). So, we set out to study the RAB14-dependent trafficking of MET. Rab coupling protein (RCP) is known to be an interacting partner of RAB14, and a study in the H1299 cell line reported it to interact with MET (45,46). RCP is also reported to be overexpressed in several cancer type and associated with metastatic property (47,48). To explore the alterations of RCP in breast cancer, we analyzed 32 publicly available breast cancer datasets consisting of a total of 15404 patients in cBioportal and found that RCP is amplified in these datasets, supporting Zhang et al (Fig. S4A) (47).

First, we analyzed the colocalization of MET and GFP-RCP with EEA1. Cells transfected with GFP-RCP were treated with HGF and then fixed, followed by immunostaining for MET and EEA1. In colocalization analysis, we found that 37 % of MET-positive RCP vesicles colocalize with EEA1 (Fig. S4B). To study whether RCP has any role in the RAB14-mediated MET recycling, we examined its subcellular localization on RAB14-MET vesicles in the presence or absence of HGF. Due to the unavailability of a validated antibody for immunocytochemistry for RAB14 and RCP, overexpression constructs were used. The anti-RCP antibody used for immunoblotting was not applicable for immunofluorescence. GFP-RCP and Cherry-RAB14 co-expressing cells were fixed and immunostained for MET. Images were captured using a confocal microscope, and the colocalization of MET-RAB14 with RCP was analyzed. In agreement with Gundry et.al, we observed an increased localization of RCP with RAB14 (35% to 51%) in the presence of HGF stimulation (46). Further, the colocalization of RCP with MET (32% to 48%) and MET-RAB14 (22% to 37%) was found to be increased upon HGF stimulation (Fig. 5A).

**Figure 5:**
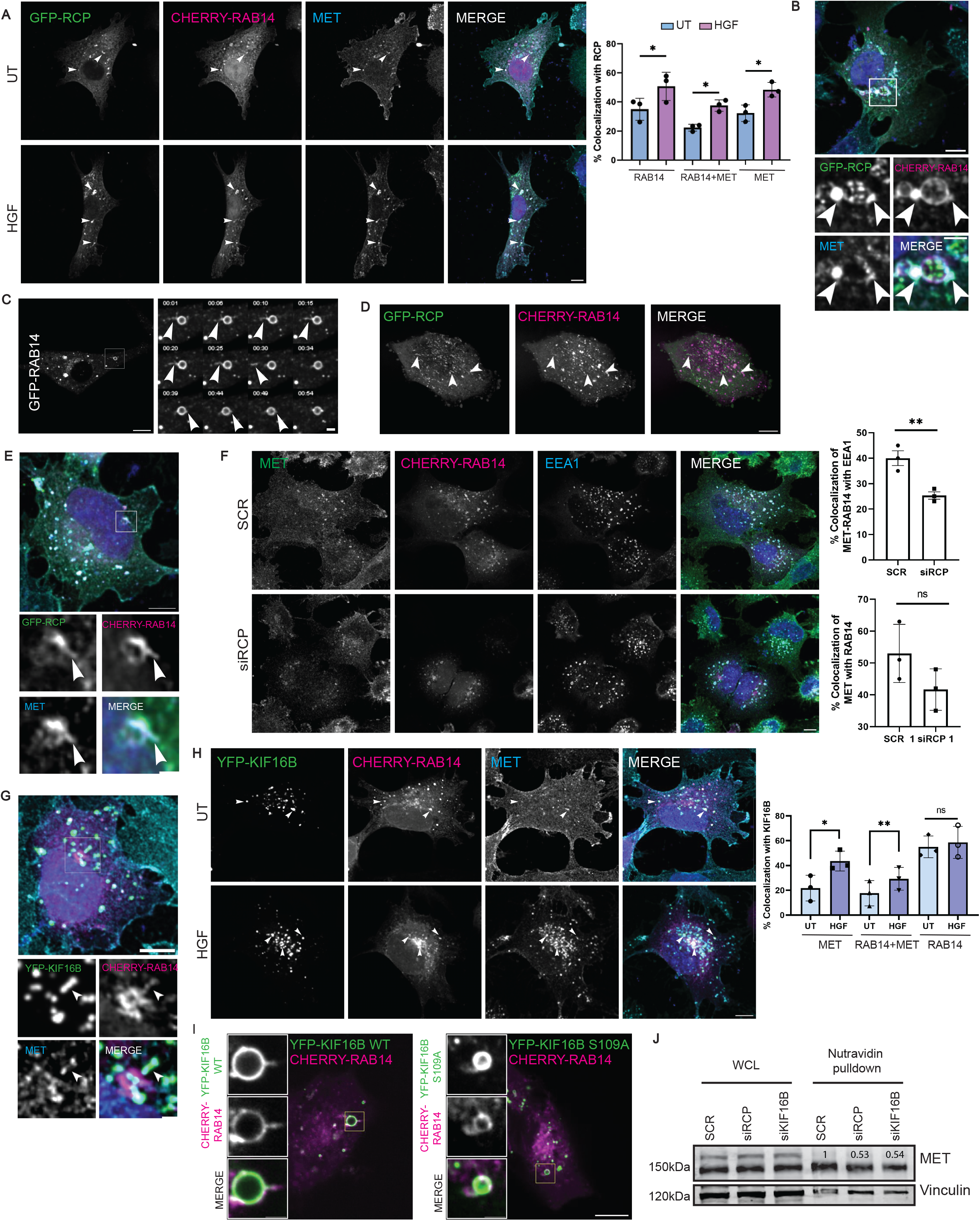
RCP-RAB14-KIF16B trafficking axis regulates MET recycling. (A) BT-549 cells seeded on glass coverslips were transfected with GFP-RCP and Cherry-RAB14. After 14h of transfection, cells were incubated for 2h with or without HGF, then fixed and immunostained for MET. Images were captured using a confocal microscope. scale bar: 10µm. The percent colocalization of GFP-RCP with RAB14 or RAB14-MET in the presence or absence of HGF treatment was calculated and plotted. scale bar: 10µm. The error bar represents Mean ±SD. Paired t-test. * P< 0.05, N=3, n=200. (B) BT-549 cells seeded on glass coverslips were transfected with GFP-RCP and Cherry-RAB14. After 14h of transfection, cells were incubated for 2h with HGF stimulation, then fixed and immunostained for MET. Images were captured using a confocal microscope. The inset shows a magnified view of the region indicated by the box. scale bar: 10µm. Inset: 2µm. (C) BT-549 cells were transfected with GFP-RAB14 and seeded on a glass-bottom imaging dish. Time-lapse images were captured using a confocal microscope. The inset shows the events over time, where the tubule is formed and, upon fission, dissociates from the vesicular body. Scalebar: 10µm, Inset: 2µm (D) BT-549 cells co-transfected with GFP-RCP and Cherry-RAB14 were seeded on a glass-bottom imaging. Time-lapse movies were captured using a confocal microscope. Arrowheads indicate the endosomal tubules containing RCP and RAB14. Scalebar: 10µm. (E) BT-549 cells seeded on glass coverslips were transfected with GFP-RCP and Cherry-RAB14. After 14h of transfection, cells were incubated for 2h with HGF stimulation, then fixed and immunostained for MET. Images were captured using a confocal microscope. The inset shows a magnified view of the region indicated by the box. scale bar: 10µm, Inset: 2µm. (F) BT-549 cells were treated with control or SMARTpool siRNA against RCP. 60h after transfection, cells were transfected with Cherry-RAB14. After providing 2 hours of HGF stimulation, cells were fixed and immunostained for MET and EEA1. Images were captured using a confocal microscope. The percentage colocalization of RAB14 endosomes containing MET with EEA1 was calculated. scale bar: 10µm. The error bar represents Mean ±SD. Paired t-test. ** P< 0.001, ns P > 0.05, N=3, n=200. (G) BT-549 cells seeded on glass coverslips were transfected with YFP-KIF16B and Cherry-RAB14. After 14h of transfection, cells were incubated for 2h with HGF stimulation, then fixed and immunostained for MET. Images were captured using a confocal microscope. The inset shows a magnified view of the region indicated by the box. scale bar: 10µm, inset: 2µm. (H) BT-549 cells seeded on glass coverslips were transfected with YFP-KIF16B and Cherry-RAB14. After 14h of transfection, cells were incubated for 2h with or without HGF, then fixed and immunostained for MET. Images were captured using a confocal microscope. scale bar: 10µm. The percent colocalization of YFP-KIF16B with RAB14 or RAB14-MET in the presence or absence of HGF treatment was calculated and plotted. scale bar: 10µm. The error bar represents Mean ±SD. Unpaired t-test. ** P< 0.01, * P< 0.05, ns P > 0.05, N=3, n=200. (I) BT-549 cells co-transfected with YFP-KIF16B WT or S109A mutant and Cherry-RAB14 were seeded on a glass-bottom imaging dish. Time-lapse images were captured using a confocal microscope. The inset shows a magnified view of the region indicated by the box. scale bar: 10µm, Inset: 2µm. N=3, n=20. (J) RCP or KIF16B were silenced using SMARTpool siRNA in MDA-MB-231 cells. After 72h of transfection of siRNA, cells were treated with HGF for 30 minutes and proceeded for surface biotinylation. The cells were incubated with biotin at 4°C for 45 minutes. Washes were given to remove excess biotin; subsequently, biotin was quenched, and cells were lysed. The lysate was allowed to bind with Neutravidin beads and the biotinylated proteins were precipitated. The samples were separated by SDS-PAGE and processed for immunoblotting. The number indicates the MET signal normalized with MET signal intensity of the respective WCL and further normalized with vinculin signal intensity.

Gundry et al demonstrated that HGF stimulation induces phosphorylation of RCP at Ser435, which enhances its complex formation with RAB14. We analyzed the distribution of MET and RAB14 in the presence or absence of HGF stimulation with the RCP S435A mutant. An increased colocalization of RCP S435A with MET (38% to 58%) and MET-RAB14 (28% to 48%) vesicles was observed upon HGF stimulation (Fig. S4C). However, the colocalization of RCP S435A with RAB14 did not increase with HGF treatment.

Interestingly, we observed that MET and RCP are enriched at the vesicular buds on RAB14 vesicles (Fig. 5B), which are critical for cargo sorting and recycling. Live cell imaging of GFP-RAB14 revealed that these buds elongate and undergo fission to form tubulovesicular structures (Fig. 5C, MOVIE 4). Live-cell confocal imaging of cells expressing GFP-RCP and Cherry-RAB14 revealed the presence of RCP in the RAB14 tubules (Fig. 5D, MOVIE 5). Subsequently, fixed cell confocal imaging highlighted that MET is present in these tubular elongations emanating from endosomes along with RCP and RAB14 (Fig. 5E). Further, the MET-containing GFP-RCP vesicles were positive for WASH staining, which promotes actin polymerization at endosomes (Fig. S4D) (28).

To investigate whether RAB14 plays a role in RCP recruitment to MET-endosomes, we overexpressed GFP-RCP in RAB14-depleted cells, then fixed and immunostained them with the anti-MET antibody. Confocal imaging revealed that the percentage of colocalization of MET with RCP was unaltered in the RAB14-silenced cells (Fig. S4E). Next, we examined the role of RCP in RAB14 and MET endosomal association. RCP was depleted with siRNA and validated with qRT-PCR and immunoblotting (Fig. S4F, G). The cells were transfected with Cherry-RAB14, treated with HGF, and fixed. Subsequently, the cells were immunostained with antibodies against MET and EEA1. Images were captured with a confocal microscope, and colocalization was quantified. A 15% decrease in colocalization of MET to RAB14 endosomes that was present with EEA1 was observed in the RCP-silenced cells compared to the control (Fig. 5F). Further, upon RCP depletion, the colocalization of MET to RAB14 was found to be reduced by 10%, but was statistically non-significant (Fig. 5F). The pool of MET or RAB14 vesicles that do not colocalize with EEA1 as observed in (Fig S3H) was unperturbed upon RCP silencing (Fig. S4I).

Molecular motors are one of the effectors of RAB GTPase, which cause membrane tubulation, thereby promoting protein sorting and trafficking. KIF16B is one of the motor proteins known to work with RAB14 (49). Hence, we analysed the colocalization of MET-containing RAB14 vesicles with KIF16B. Cells expressing YFP-KIF16B and Cherry-RAB14 were fixed and immunostained for MET. Confocal imaging revealed that KIF16B also resides in the endosomal tubules along with RAB14 and MET (Fig. 5G). Next, we analyzed the colocalization of RAB14-MET endosomes with KIF16B in the presence or absence of HGF. YFP-KIF16B and Cherry-RAB14 co-expressing cells were fixed and immunostained for MET. Images were captured using a confocal microscope, and percentage colocalization was quantified using Motiontracking. An increased colocalization of MET (21% to 43%) and MET-RAB14 (17% to 29%) vesicles with KIF16B was observed in the presence of HGF (Fig. 5H). However, RAB14 colocalization to KIF16B remained unaltered upon HGF treatment (Fig. 5H). Since KIF16B is an effector of RAB14, upon silencing of the GTPase, the colocalization of KIF16B with MET was found to be reduced, as expected (Fig. S4J) (49).

KIF16B is known to promote tubulation in the early endosomes (50). So, to investigate the potential role of KIF16B in RAB14 endosomal tubulations, we overexpressed WT or the dominant negative form of the motor with an Ala substitution at the S109 position of KIF16B. BT-549 cells co-expressing Cherry-RAB14 with WT KIF16B or the S109A mutant were subjected to live cell imaging. In 40% of the WT KIF16B expressing cells, RAB14 tubules were observed, whereas it was only 15% in the case of the S109A mutant expressing cells (Fig. 5I, MOVIE 6). This result suggests a potential role for KIF16B in endosomal tubule formation.

Next, to investigate the role of RCP or KIF16B in the surface delivery of MET, we quantified total MET pool at the surface employing surface biotinylation. We performed the surface biotinylation in MDA-MB-231, since a substantial loss of cells was observed in BT-549 during the experimental procedures resulting in insufficient cell lysate for downstream analysis. The RCP or KIF16B silenced cells were treated with HGF for 30 minutes and proceeded for surface biotinylation. Vinculin is taken as a loading control as it is reported to also present in the membrane fraction (51). The MET population at the cell surface was found to be reduced upon RCP or KIF16B silencing, indicating their role in the delivery of MET to the plasma membrane (Fig 5J, S6). Depletion of RCP or KIF16B did not alter the MET total protein expression (Fig. S4H, S6).

Next, we performed gelatin degradation assay to investigate the role of RAB14, RCP or KIF16B in TNBC invasion. MDA-MB-231 or BT-549 cells were transfected with SMARTpool siRNA targeting the respective genes or control siRNA, subsequently seeded on labelled gelatin and the degradation index was analyzed. We observed a significant reduction in the gelatin degradation in MDA-MB-231 as well as BT-549 cells upon silencing of RAB14, RCP or KIF16B compared to control (Fig S4K-L). To validate this observation, we used single oligoes targeting RAB14 and RCP and performed gelatin degradation assays. Consistent with the SMARTpool siRNA data, silencing of RAB14 or RCP using individual oligoes also showed a reduction in the gelatin degradation activity (Fig S4L).

Together, these observations suggest that RAB14-RCP-KIF16B axis facilitates the MET surface delivery through tubulovesicular carriers.

### HGF-MET facilitates ECM degradation by recycling MT1-MMP to the surface

MT1-MMP is one of the invadopodia-associated surface proteases with a wide range of substrate specificity for diverse ECM components (52). Perturbed MT1-MMP recycling impedes ECM degradation and cancer invasion (6,53). HGF is known to increase the expression and activity of invadopodia-associated metalloproteases MMP2 &MMP9 (54). We asked if HGF-stimulated invadopodia activity is mediated by MT1-MMP. We began by studying the expression of MT1-MMP in HGF-treated conditions. 3h of HGF stimulation did not alter the expression or degradation of MT1-MMP (Fig. S5A, A’, S6).

Next, we investigated whether HGF treatment leads to enhanced recycling of the protease. We performed an antibody uptake assay, where cells were incubated with MT1-MMP antibody at 4, followed by internalization at 37□. Antibodies available at the surface were removed, and cells were incubated with HGF to allow recycling at 37□. The percent of recycling was calculated as described in the methods. Antibody uptake assay revealed that HGF stimulation induced 30% recycling of antibody-bound MT1-MMP, whereas, in the untreated condition, the recycling of MT1-MMP was 10% (Fig. 6A).

**Figure 6:**
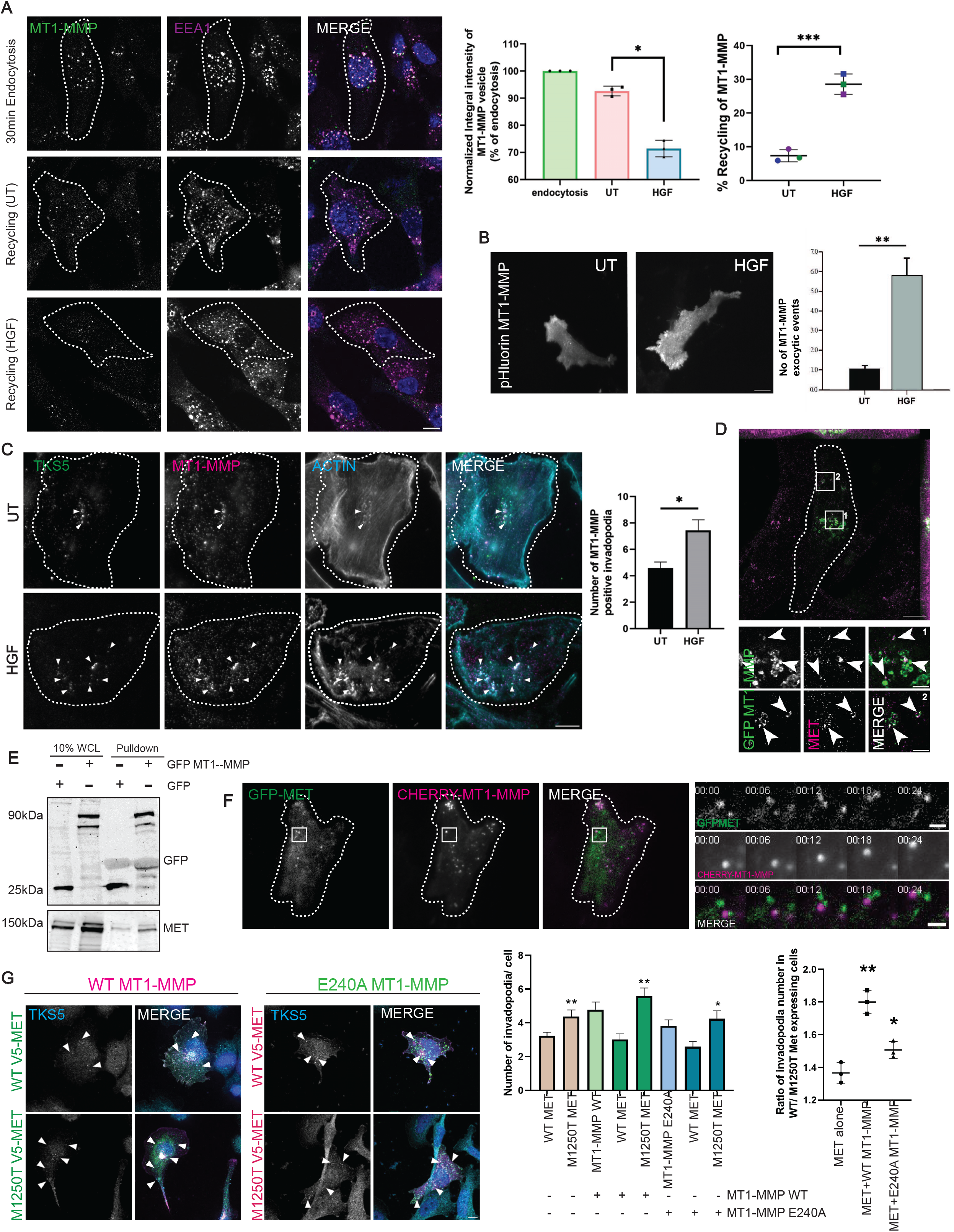
HGF/MET facilitates ECM degradation by recycling MT1-MMP to the surface. (A) MDA-MB-231 cells seeded on coverslips, surface labeled with MT1-MMP antibody at 4°C for 1 h Complete media was added, and cells were shifted to 37°C for 30 minutes to allow endocytosis. To remove surface-bound antibodies, an acid wash was given. After washing with PBS, serum-free media was added with or without HGF, and the cells were shifted to 37°C for 10 minutes to allow recycling from the endocytosed pool. Cells were fixed and immunostained for EEA1. The integral intensity of the MT1-MMP and the percentage of MT1-MMP antibody recycling were calculated. N = 3, n= 300; scale bars = 10 µm. The error bar represents means ± SD. Paired t-test, *** P< 0.001, Unpaired t-test, * P <0.05. (B) MDA-MB-231 cells transfected with pHluorin MT1-MMP were seeded on gelatin-coated glass-bottom dishes. Time-lapse imaging was performed in the presence or absence of HGF stimulation using a Nikon TIRF microscope without any intervals for 300 frames. The flashes of pHluorin MT1-MMP, which represent exocytic events, were calculated using Motiontracking. scale bars = 10 µm. The error bar represents Means ± SEM. Unpaired t-test, ** P < 0.001. N = 4, n= 35. (C) MDA-MB-231 cells were seeded on gelatin-coated glass-bottom dishes and allowed to attach for 3h, followed by incubation with or without HGF for another 3h. Cells were fixed and immunostained for TKS5, MT1-MMP, and F-actin. Images were captured using a Nikon TIRF microscope. TKS5, MT1-MMP, and Actin-positive structures were identified using Motiontracking. The error bar represents Mean ±SEM, Unpaired t-test, * P < 0.01. scale bar: 10µm. N=3, n=120. (D) GFP-MT1-MMP expressing MDA-MB-231 cells were fixed and immunostained for MET. Images were captured using an LSM900 Airyscan super-resolution microscope. Images were processed using the Airyscan Joint deconvolution (jDCV) method. The image was converted to Maximum Intensity projection for representation. The inset shows a magnified view of the region indicated by the box. Scale bar: 10 µm, inset: 2 µm. (E) GFP or GFP-MT1-MMP expressing BT-549 cells were lysed and incubated with GST agarose beads bound to 25µg of GFP binding protein (GBP). Next washes were given to remove unbound proteins and proceeded for immunoblotting with GFP and MET. WCL: whole-cell lysates (F) BT-549 cells co-expressing GFP-MET and Cherry MT1-MMP were subjected to live-cell TIRF imaging. The inset represents an event at different time points. Scale bar: 10 µm, Inset: 2 µm. (G) BT-549 cells expressing MT1-MMP WT or a catalytic mutant alone or with V5-MET WT or M1250T mutant were seeded on gelatin-coated coverlips for 6h and fixed. Cells were immunostained for V5 and TKS5. Images were captured with the confocal microscope. The number of TKS5 puncta in the MT1-MMP and/or MET expressing cells was calculated and plotted. The error bar represents Mean ±SD, Unpaired t-test. scale bar: 10µm. ** P<0.01, * P < 0.05, N=3, n=150.

To support our observation from the antibody uptake assay, we employed TIRFM-based imaging to capture MT1-MMP exocytosis in real time. We overexpressed a pH-sensitive GFP, pHluorin-tagged MT1-MMP in MDA-MB-231, and quantified the number of exocytic events using TIRFM. We could see an increased surface delivery of pHluorin MT1-MMP when stimulated with HGF (Fig. 6B, MOVIE 7A, B).

The observation was further validated using surface biotinylation. For biotinylation, untreated or HGF-treated cells were incubated with uncleavable biotin, EZ-linked Sulfo-NHS biotin at 4□. The cells were lysed, and the biotinylated pool was captured with streptavidin beads. Immunoblot analysis revealed that the HGF-treated cells have a higher surface MT1-MMP pool than the untreated cells (Fig. S5B). CIMPR has been taken as control, whose surface level did not alter with HGF treatment (Fig. S5B’).

Next, we asked whether HGF also promotes the endocytosis of MT1-MMP. To address this, an antibody uptake assay was performed in the presence or absence of HGF. Cells grown on coverslips were incubated with MT1-MMP antibody at 4 for 1h. The cells are allowed to internalize the antibody in the presence or absence of HGF. Cells were fixed, imaged with a confocal microscope, and the integral intensity of the internalized MT1-MMP antibody was quantified. HGF stimulation did not show any effect on the endocytosis of MT1-MMP (Fig. S5C).

MT1-MMP is recruited to invadopodia for efficient ECM remodelling. Since we saw an increased MT1-MMP recycling to the surface, we investigated the MT1-MMP recruitment to invadopodia in the presence or absence of HGF. MDA-MB-231 was seeded on gelatin with or without HGF. After 6h, cells were fixed and immunostained for invadopodia markers. TIRF imaging revealed that, indeed, HGF treatment leads to more MT1-MMP-containing invadopodia compared to the untreated condition (Fig. 6C). Further, to understand whether the receptor MET is also involved in the MT1-MMP recruitment to invadopodia, MET was silenced using siRNA, and the MT1-MMP population at invadopodia was quantified. Upon depletion of MET, the number of MT1-MMP-containing invadopodia was reduced (Fig. S5D). However, MET depletion did not alter MT1-MMP expression (Fig. S1I).

Next, to gain insight into the possible role of MET in MT1-MMP recruitment to invadopodia, we examined the localization of MT1-MMP with MET. The percentage of colocalization was increased from 12% to 45% when the cells were stimulated with HGF (Fig. S5F). Super-resolution microscopy revealed that in GFP-MT1-MMP expressing cells, MET localizes to the same endosomal microdomain as MT1-MMP (Fig. 6D). Interestingly, a protein-protein interaction prediction database PSOPIA anticipated the interaction between MT1-MMP and MET (55). To establish the physical interaction between MET and MT1-MMP, we employed a GFP nanobody pulldown approach. Lysates of GFP or GFP-MT1-MMP expressing cells were incubated with GST bead-bound GFP binding protein. The interaction was analyzed by immunoblotting. An increased binding of MET to GFP-MT1-MMP compared to GFP was observed (Fig. 6E, S6). The interaction was supported by performing GFP nanobody pulldown in HeLa and MCF10A DCIS (Fig. S5F). Next, to validate the interaction, we performed a reverse pull-down. We overexpressed GFP or GFP-MT1-MMP with V5-MET. V5-MET was trapped using Ni-NTA beads. Immunoblotting of the pulldown sample revealed that V5-MET exclusively interacts with GFP-MT1-MMP (Fig. S5G, S6). However, immunoprecipitation with endogenous MET we observed a weak interaction of MT1-MMP with MET (Fig S5H, S6).

MT1-MMP is known to cleave and shed the ectodomain of LDL receptor, apo-A1, ApoE, p-selectin, MMP2, MMP13, and αV integrin (56–59). As MT1-MMP and MET interact, we asked whether MT1-MMP can also activate MET by shedding its pro-domain. We quantified the mature MET population of 140 kDa in MT1-MMP-silenced cells by immunoblotting. The total matured MET population remains unchanged in the protease-depleted cells, whereas GFP-MMP2 which act as a substrate for MT1-MMP showed increase in the protein level (Fig. S5I, S6). Further, to comprehend the relevance of the interaction between the RTK and MT1-MMP, we hypothesize that their physical association plays a crucial role in their simultaneous enrichment at invadopodia, potentially via co-trafficking. To test this hypothesis, cells were transfected with GFP-MET and Cherry-MT1-MMP and imaged under a TIRF microscope. In live-cell TIRFM, GFP-MET and Cherry-MT1-MMP were found to be co-transported at the cell surface (Fig. 6F, MOVIE 8). These data suggest that HGF promotes MT1-MMP surface delivery and the receptor MET via physical interaction with MT1-MMP, ensuring a directed delivery to invadopodia.

A study by Ferrari et.al. suggests that MT1-MMP is not only required for invadopodia maturation but also can affect invadopodia formation (60). To explore the role of MT1-MMP on invadopodia formation, the protease was overexpressed, and the number of invadopodia formed was quantified. The overexpression of MT1-MMP WT or a catalytically inactive mutant E240A MT1-MMP increases the invadopodia number; however, no significant change was observed between the E240A MT1-MMP and WT MT1-MMP (Fig. S5L, L’). Next, we analyzed the number of invadopodia in the MT1-MMP knocked-out cells. Interestingly, the invadopodia formation was drastically reduced in the MT1-MMP-deleted cells compared to the control, highlighting the significance of MT1-MMP in invadopodia formation (Fig. S5M).

Interestingly, when we co-expressed MT1-MMP and MET together, we observed an increased invadopodia formation. The co-expression of MET WT or M1250T with WT MT1-MMP leads to a 1.8-fold increase in the difference observed in the number of invadopodia formed between the WT and M1250T MET compared to the 1.35-fold increase in overexpression of MET WT and M1250T alone (Fig. 6G). However, E240A MT1-MMP could enhance invadopodia formation up to 1.5-fold in M120T MET expressing cells (Fig. 6G). This highlights the potential influence of MT1-MMP on invadopodia formation. When we analyzed the MT1-MMP recruitment to invadopodia in the presence of the M1250T mutant, we observed an increased colocalization of MT1-MMP to invadopodia compared to the WT MET (Fig. S5K).

Further, to understand whether the increased invadopodia number in the M1250T MET and MT1-MMP (Fig. 6G) was due to the higher affinity of the MET mutant for MT1-MMP, we employed a pulldown approach. Cell lysates from V5/His-MET WT or M1250T and GFP or GFP-MT1-MMP were incubated with Ni-NTA beads. The pulldown samples were analyzed by immunoblotting. We did not observe a preferential binding of MT1-MMP to M1250T (Fig. S5N, S6), nor was the MT1-MMP expression altered in the mutant expressing cells (Fig. S5J, S6), inferring the increased invadopodia formation of M1250T MET in the presence of MT1-MMP is not because of physical interaction.

Taken together, our observations suggest that the HGF-MET axis directs MT1-MMP to invadopodia. The physical association between MT1-MMP and RTK MET ensures their co-trafficking to the surface, thereby promoting invadopodia formation.

## DISCUSSION

The current study elucidates how intracellular trafficking of MET could promote invadopodia-driven breast cancer invasion in TNBC cells. The results highlight- (i) HGF stimulation promotes MET and MT1-MMP recycling to invadopodia, (ii) MET and MT1-MMP physically interact, which we believe to be crucial for their co-trafficking to the cell surface, (iii) RAB4 and RAB14 facilitates the MET delivery to invadopodia, (iv) The RTK colocalizes with RCP on RAB14 endosomal tubules, (v) the formation of which is promoted by KIF16B for efficient MET trafficking.

RTK signalling was shown to play a critical role in invadopodia formation by coordinating activation of key invadopodia associated proteins and subsequent actin remodelling (61). In the current study we have demonstrated that MET or EGFR signalling facilitate invadopodia formation with the RTKs getting enriched at invadopodia in a ligand dependent manner (Fig. 1A-F, 2A-G, S2H). This enhanced recruitment of RTKs to invadopodia is potentially due to targeted delivery of RTKs through increased recycling upon growth factor stimulation. Consistent with our observation, a previous study have reported that SNARE-mediated delivery of EGFR to invadopodia facilitates invasive activity (11). Together, this suggests that growth factor-mediated recruitment of RTKs to invadopodia may be a generalized mechanism applicable to RTKs that are known to promote invadopodia-associated activities.

Further, our study reveals that MET localizes only to a sub-set of TKS5-positive invadopodia. The mature degradation-positive structures largely devoid of the RTK, suggesting a temporal association of MET with invadopodia at an early stage of the biogenesis (Fig. 2D, F). TKS5 is one of the crucial scaffold proteins required during invadopodia formation and maturation. Kreider-Letterman et al have shown that accumulation of TKS5 gradually increases as invadopodia mature (37). We observed an inverse correlation between MET and TKS5 levels at the TKS5 enriched invadopodia (Fig. 2E). Analysis of the MET/TKS5 fluorescence intensity ratio demonstrated that the majority of invadopodia exhibited a low ratio (Fig. S2B). Together, these observations suggest that MET is preferentially associated with nascent structure and may function during the early stages of invadopodia biogenesis prior to maturation, which is marked by progressive TKS5 recruitment (62). Further investigation of MET localization to invadopodia using live cell time-lapsed microscopy would enable us to analyze the temporal dynamics of MET recruitment to invadopodia and establish its temporal role during the biogenesis process.

It is well established that HGF treatment leads to endocytosis of the RTK leading to its degradation in lysosomal compartments (63). In the current study we also observed a pool of RTK that is recycled to invadopodia upon HGF stimulation (Fig 2). Our results further suggested that recycling of the RTK is associated with invadopodia mediated ECM degradation (Fig 3, S2). These observations corroborated well with enhanced invasive phenotype exhibited by the TNBC cells that overexpressed degradation defective mutant of MET which possess enhanced recycling ability (Fig. 3E) (31). However, the endocytosis defective mutant failed to promote invasive behaviour to a similar extend, suggesting that receptor internalization with subsequent surface recycling is critical to drive the invasive activity. Previous reports demonstrated that in response to HGF stimulation, GGA3 and CLIP170 regulate RAB4-mediated recycling of MET driving cancer cell migration (64,65). Here, we identified a previously unknown, RAB14-mediated recycling pathway of MET, that promotes invadopodia formation and TNBC invasion. Moreover, the MET localized to distinct endosomes positive for RAB4 or for RAB14 near the cell cortex, implicating for discrete recycling pathways regulated by these RABs (Fig. S3I). RAB14 is associated with the recycling of cargoes like MT1-MMP, which is one of the most well-studied invadopodia-associated metalloproteases (66). Perturbation in the delivery of MT1-MMP to invadopodia impairs ECM degradation and invasion (6,7). Our observations suggest that the HGF-MET axis promotes co-trafficking of MT1-MMP and the RTK to invadopodia, potentially facilitated by their physical interaction as demonstrated by immunoprecipitation assay (Fig. 6).

The role of MT1-MMP during invadopodia maturation is well established (67,68). The protease facilitates cell invasion via its degradative activity on ECM components. Interestingly, recent literature reported an emerging role for the protease during the initial stage of invadopodia formation (60,69). In agreement with these reports, overexpression or depletion of MT1-MMP altered invadopodia formation (Fig. S5L-M). Additionally, when MT1-MMP was co-expressed with the WT or the M1250T MET, the mutant expressing cells showed enhanced invadopodia formation compared to WT MET (Fig. 6G, S5L). The effect of MT1-MMP co-expression on invadopodia formation was also exhibited by the catalytically inactive mutant. Collectively, these findings highlight the role of MT1-MMP in invadopodia formation that is independent of its catalytic function. We speculate that the increased recycling ability of the MET mutant may promote the targeted delivery of MT1-MMP to invadopodia through their physical association during the initial stages of invadopodia biogenesis.

### Regulation of MET trafficking through RCP-RAB14-KIF16B axis

RAB GTPases orchestrate cargo trafficking by recruiting diverse sets downstream molecules that operate on the sorting and the delivery of the cargo to destined membrane (22). Adapter proteins represent a class of the RAB-effectors that helps in efficient cargo trafficking by bridging the cargo and the trafficking machinery (70). Among these, RAB11 family- interacting proteins (RAB11FIPs) are RAB11 binding proteins that engage endocytic vesicles with motors (70). RCP, a member of the RAB11FIP family, interacts with multiple RAB GTPases, including RAB11, RAB4, and RAB14, as well as cargos such as MET (45,71,72). Gundry et al have established that RAB14 along with RCP co-ordinates HGF dependent trafficking of Epinephrin receptor2 (EphA2) (46). This study further demonstrated that the HGF mediated phosphorylation of RCP at S435 strengthens its binding with RAB14. We showed unaltered colocalization of RAB14 with RCP S435A in response to HGF, which substantiate the findings of Gundry et al (Fig S4C). However, our colocalization analysis showed that HGF stimulation enhances the association of MET with both WT and phosphodeficient RCP mutant, suggesting this HGF-induced RCP phosphorylation is not crucial for its association with MET (Fig S4C). Notably, the reduced colocalization of MET and RAB14 at the EEA1 compartment upon silencing of RCP indicate that RCP may function as a bridging molecule between RAB14 and MET facilitating efficient MET recycling (Fig. 5F) (45,46).

RAB14-positive endosomes are known to generate bud-like microdomains that function as specialized hubs for selective cargo sorting (44). These bud like specialized microdomains serve as a platform for cargo segregation through recruitment of specific trafficking machineries that direct proteins towards recycling pathway (73). In our study, we observed that these RAB14 buds elongate to tubular structures containing both MET and RCP (Fig. 5B-E, MOVIE 5). Moreover, KIF16B was also found to be present on these tubular vesicles (Fig 5G). KIF16B is an established downstream effector of RAB14 that facilitates the vesicle movement containing cargo like MT1-MMP and FGFR (49,66). Our findings further showed that KIF16B promotes the surface delivery of MET by promoting RAB14-positive tubulovesicular structures (Fig. J, MOVIE 6). Previous studies have demonstrated that molecular motors play important roles not only in vesicular transport but also in endosomal tubule generation, with KIF16B promotes the endosomal tubulation in early endosomes, while KIF13A promotes tubulation in RAB11 recycling endosomes (52,76). Collectively, these studies together with our findings demonstrate the crucial role of molecular motors in regulating membrane tubulation and driving distinct trafficking pathways.

Together our study provides a mechanistic understanding of the HGF-MET-mediated invadopodia formation and MT1-MMP recycling (Fig. 7). We present evidence demonstrating MET localization to invadopodia is regulated by RAB4 and RAB14- dependent recycling in response to HGF stimulation. We speculate that RCP forms a trafficking-assembly with MET and RAB14 on the sorting endosomes. This trafficking- assembly localizes to the endosomal buds, which in turn elongates due to the activity of KIF16B forming tubular carriers. Thus, the RCP co-ordinated association of MET with RAB14-KIF16B may be a possible mechanism underlying the sorting of the RTK from the endosomal subdomains directing its trafficking to the cell surface. In addition, functional interaction between MET and MT1-MMP facilitates their co-transport to invadopodia, driving ECM degradation during TNBC invasion.

**Figure 7:**
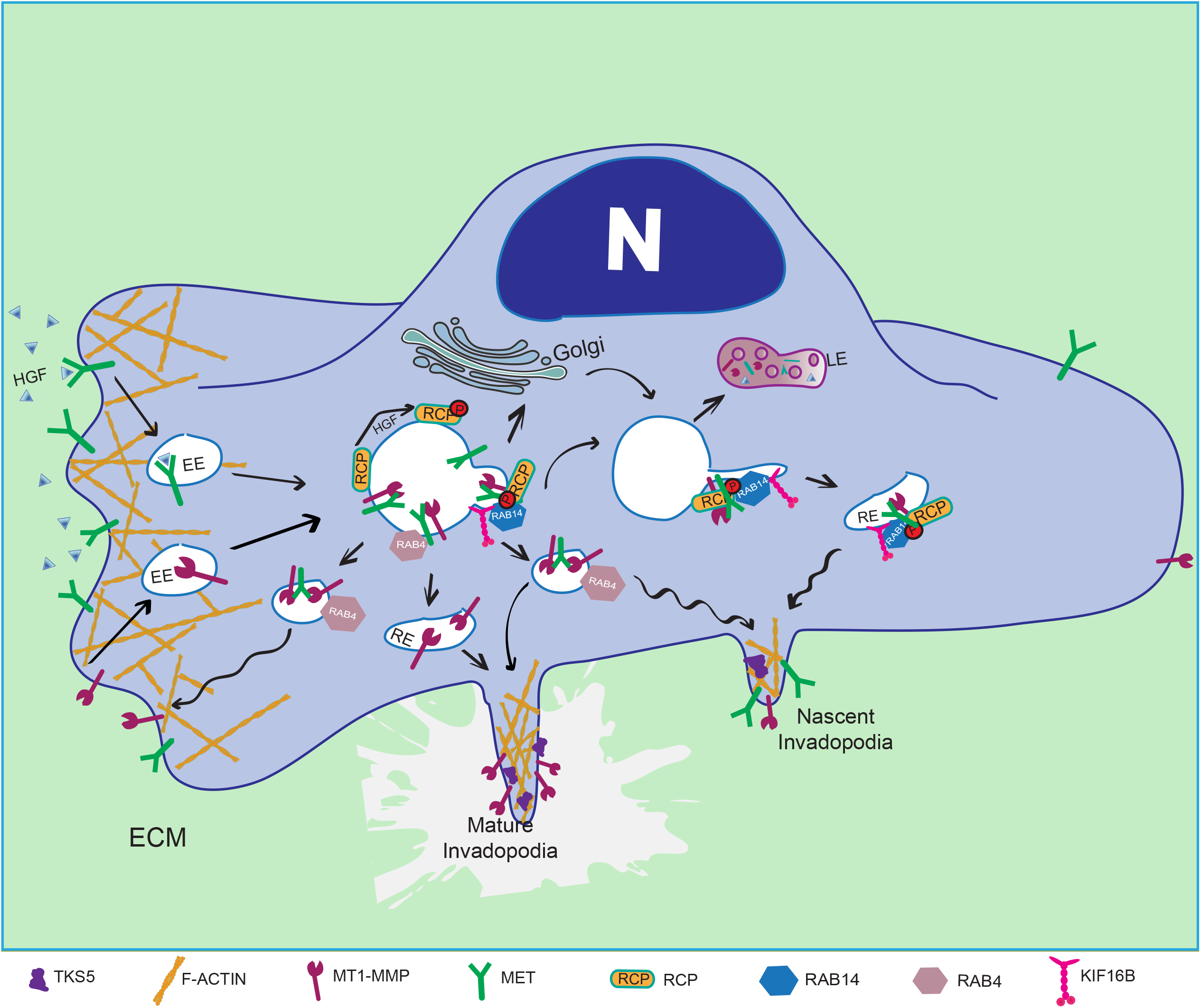
**Proposed model showing RAB14-mediated MET trafficking to invadopodia along with MT1-MMP in the presence of HGF stimulation**. HGF stimulation triggers invadopodia formation and ECM degradation by promoting invadopodia formation and MT1-MMP recycling. HGF also increases the recruitment of MET to invadopodia by stimulating the trafficking of the RTK in a RAB4 or RAB14-dependent trafficking pathway. HGF stimulation increases the phosphorylation (46) and colocalization of RCP with MET-containing RAB14 vesicles. RCP and MET are present on the endosomal budding site of RAB14, which further elongate and form tubules. KIF16B gets recruited to these endosomes through RAB14 and drives the endosomal tubulation for efficient cargo trafficking. MT1-MMP interacts with MET, presumably promoting their co-trafficking to invadopodia.

Although the role of HGF/MET in tumor progression is well established for other cancer types, to investigate whether non-TNBC subtypes also adopt the mechanism identified in the present study to facilitate cancer invasion requires extensive experimental evidence.

## Material and Methods

### Plasmids, antibodies, and reagents

The following antibodies were purchased commercially: rabbit anti-FISH (TKS5) (SantaCruz, sc-30122), 1:500 (IF), mouse anti-cortactin (Millipore, 05–180), 1:300 (IF); (SantaCruz, sc-55579) 1:500 (IF), rabbit anti-MET (CST, D1C2 XP) 1:400 (IF), 1:1000 (IB), mouse anti-MET (CST, L6E7) 1:500 (IF), 1:1000 (IB), rabbit anti-Phospho-Met (Tyr1234/1235) (CST, D26 XP) 1:800 (IF), 1:1000 (IB), mouse anti-V5 (Sigma, SV5-Pk1) 1:400 (IF), 1:1000 (IB), rabbit anti-Rab11Fip1 (Protein technology, 16778-1-AP) 1:1000 (IB), rabbit anti-mouse MT1-MMP (Millipore, MAB3328), 1:1,000 (IB) and 1:400 (IF); mouse anti-MT1-MMP (R&D Systems, MAB9181-SP), 1:200 (IF), mouse anti- Vps35(SantaCruz, sc-374372) 1:300 (IF); rabbit anti-Vps26 (Abcam, ab23892) 1:1000 (IB), rabbit anti-EGFR (CST, D38B1), rabbit anti-Golgin97 (CST, D8P2K) 1: 200 (IF); mouse anti-vinculin (Sigma, V9131), 1:1,000 (IB); rabbit anti-actin (Sigma, A2066), 1:1,000 (IB); mouse anti-γ-tubulin (Sigma, T6557), 1:3000 (IB), mouse anti-GFP (Roche, 11814460001), 1:2000 (IB); mouse anti-Rab5 (BD Biosciences, 610724), 1:300 (IF); mouse anti-His (ThermoFischer Scientific), 1:1,000 (IB), Rabbit anti-MMP2 antibody (Proteintech, 10373-2- AP), 1:1000 (IB). The anti-rabbit EEA1 antibody was a kind gift from Professor Marino Zerial (Max Planck Institute of Molecular Cell Biology and Genetics, Dresden, Germany) and was used at 1:1000 (IF). Rabbit anti-WASH antibody was kindly shared by Professor Alexis Gautreau, used at 1:300 (IF).

pLenti MET-GFP (37560) and pCDNA3.1 MET-V5/His (201982) were purchased from Addgene. MET was amplified from pLenti MET-GFP using PCR and cloned into pEGFPN1 vectors using the XhoI and HindIII sites. M1250T and LL/AA mutants were generated in pCDNA3.1 MET-V5/His using site-directed mutagenesis. Primer details are provided in the Table 1. pLVX Tre3G MET-GFP WT was constructed by Mutagenex. pLVX TRE3G MET- GFP M1250T was generated by BIOBOX.

### Cell culture

MDA-MB-231 (HTB-26), BT-549 (HTB-122) and HeLa (CCL-2) cell lines were purchased from ATCC and verified to be free of contaminants. All the cell lines were cultured for maximum of 20 passages and then discarded. MDA-MB-231 cells were maintained in L-15 (Invitrogen) media with 10% FBS, 100 µg ml^−1^ penicillin, and 100 µg ml^−1^ streptomycin at 37°C in CO_2_-free conditions. BT-549 cells were maintained in RPMI (Invitrogen) supplemented with 0.023U/ml of Insulin, 10% FBS, 100 μg/ml penicillin, 100 μg/ml streptomycin, at 37°C with 5% CO2. Professor Phillipe Chavrier from the Centre National de la Recherche Scientifique in Paris, France, kindly shared the MCF10A DCIS cell line, which was maintained at 37°C with 5% CO2 in Advanced DMEM-F12 media supplemented with 2 mM glutamine and 5% horse serum (Invitrogen), 100 µg ml^−1^ penicillin, and 100 µg ml^−1^ streptomycin. HeLa cells were cultured using Dulbecco’s modified Eagle’s medium (DMEM) supplemented with 10% FBS, 100 μg/ml penicillin, 100 μg/ml streptomycin, and 2 mM glutamine, at 37°C with 5% CO2.

### Transient transfection and Generation of stable cell lines

50,000 cells per well were seeded on coverslips in 24-well plates one day before the experiment to transfect expression plasmids transiently. Cells were transfected with 0.5 µg of plasmid DNA using LTX/Plus transfection reagent (Life Technologies) for MDA-MB-231 cells and Lipofectamine 3000 for BT-549 cells. HeLa cells were transfected with polyethylenimine (PEI-1mg/ml), 5µl of PEI /1µg of DNA.

To generate a stable cell line, 2 × 10^6^ cells per ml were seeded in a 35 mm dish and, after 24 h, were transfected with pLVX TRE3G GFP-MET WT or M1250T with pLVX-pEF1a- Tet3G in a 4:1 ratio. Transfected cells were selected using complete media containing 800 µg/ml G418 and 1µg/ml puromycin. The media was changed every third day. Cells that survived antibiotic stress proliferated and formed colonies, were collected, and grown. Each clone was detected for GFP expression by performing immunoblotting and immunofluorescence. After antibiotic selection, cells were maintained in media containing 500 µg/ml G418 and 1µg/ml puromycin. For gene expression, 50-100 ng/ml of Doxycycline was used.

### siRNA and shRNA transfection

OntargetPlus siRNA SMARTpool targeting the gene of interest was purchased from Dharmacon. All siRNAs were used at the working concentration of 10–20 nM. Cells were seeded 24 h before performing siRNA transfection. The manufacturer’s protocol was used for transfection using Dharmafect as the transfection reagent. The sequences of SMARTpool siRNAs are provided in the Table 2.

All shRNA clones were part of the MISSION shRNA product line from Sigma-Aldrich. The TRC1.5 pLKO.1-puro non-mammalian shRNA Control Plasmid DNA (SHC002) is a negative control containing a sequence that should not target any known mammalian genes but engage with RISC. Cells were seeded 24 h before performing shRNA transfection. Cells were transfected at 50–60% confluency with 0.5 μg of shRNA plasmid DNA using LTX/Plus transfection reagent (Life Technologies). The sequences for MISSION plasmid shRNAs are provided in the Table 3.

### RNA extraction and quantitative real-time PCR

Total RNA was extracted from the cells using RNAeasy kit (QIAGEN, Cat# 74104), and cDNA was prepared using the High-Capacity RNA-to-cDNA kit (Life Technologies, Cat# 4387406). Real-time qPCR reactions were performed using the SYBR Green Kit and corresponding primers on Applied Biosystems 7300 Real-Time PCR System or Thermo Quant Studio 3.0. The primer details are provided in the Table. 1.

### Immunofluorescence and confocal imaging

Immunofluorescence was performed as described earlier by Parveen et al (53). Cells were fixed in 4% paraformaldehyde (PFA) for 15 min at room temperature followed by permeabilized with 0.1% Triton X-100 for 10 min at room temperature. The cells were blocked with 5% FBS in 1× PBS and were immunostained with the primary antibodies, followed by Alexa-labeled secondary antibodies. The coverslips were mounted using Mowiol (Calbiochem, Cat. 475904) on glass slides and imaged using an LSM780 or FV3000 confocal microscope. Data from three independent experiments were subjected to analysis by Motiontracking.

### Object-based colocalization analysis

Colocalization of objects were quantified using Motiontracking as described by Parveen et al (53). The objects were identified as vesicles in each channel depending on their size, fluorescence intensity, elongation, etc by Motiontracking. Objects identified in two different channels were considered to be colocalized if the relative overlap of respective areas was >35%. The colocalization value is the ratio of the integral intensities of colocalized objects to the integral intensities of all objects in the base channel. The values can range from 0.0 to 1.0. The random colocalization was estimated by random permutation of object localization in different channels, and the apparent colocalization was corrected for random colocalization. Similar way, the Motiontracking defines multicolor objects as those objects with a colocalization value greater than 0.35.

### CTCF calculation using ImageJ

To calculate Corrected Total cell fluorescence (CTCT) in ImageJ, a cell boundary was drawn. The area, integrated density, and mean gray value of the ROI were calculated by using the “measure” function under “Analyze”. To correct the background fluorescence, an ROI of similar size was drawn in the background where no cell is present, and the mean gray value was calculated. The corrected total cell fluorescence (CTCF) was then calculated using the following formula:

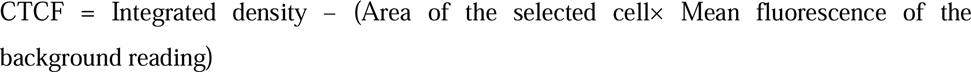

### Live-cell imaging

Cells were transfected with GFP- and/or mCherry-fused proteins alone or together for 12 h. Cells were trypsinized and seeded on glass-bottom dishes. Cells in complete medium were allowed to get attached at 37°C and further imaged with an Olympus IXplore spinSR microscope with a 100× Plan Apo N objective (oil, 1.42 NA) on an inverted stage equipped with a Tokai Hit onstage incubation system. Images were acquired and analyzed using Motiontracking.

### TIRF microscopy and analysis

#### Live-cell TIRF-M

To specifically capture events occurring near the cell surface, imaging was performed using a Nikon Eclipse Ti2-E inverted microscope equipped with an H-TIRF module and an Okolab on-stage incubation system for temperature and CO regulation. Cells expressing GFP and/or mCherry fusion proteins were seeded on gelatin-coated, glass-bottom dishes containing complete medium. Dual-color imaging was carried out simultaneously using an Apo-TIRF 60× oil DIC N2 objective lens (NA 1.49) and a Nikon LU-N4 laser combiner, with two Orca-Flash4.0 V3 sCMOS cameras (Hamamatsu Photonics). Images were acquired using NIS-Elements AR software version 5.11.00. GFP-tagged proteins were excited using a 488 nm laser, and mCherry-tagged proteins were excited using a 561nm laser.

#### Calculation of MT1-MMP exocytosis

The movies were imported into the MotionTracking software and processed as described by Goto-Silva et.al. (74). Briefly, to distinguish endosomes from background fluorescence, we applied the Bayesian Foreground/Background Discrimination (BFBD) filter, as implemented in the MotionTracking software (https://Motiontracking.mpi-cbg.de). The BFBD algorithm separates the image intensity into a fast-changing foreground and a slowly changing background, corresponding to moving vesicles and cytoplasmic fluorescence, respectively. This procedure efficiently suppresses the Poisson noise of fluorescence. Since the BFBD attributed the signal from non-moving objects to the background, we sum the previously found foreground and background. Then, a new background estimation was performed as described in Kalaizidis et al (75) with a window size of 1µm and subtracted from the de- noised images. The images were thresholded by signal-to-noise ratio (SNR) = 2.0, where the amplitude was estimated as signal uncertainty predicted by the BFBD algorithm. The resulting images were segmented by the watershed algorithm. Detected endosomes were associated with the tracks by a dynamic programming algorithm that performs local-in-time/global-in-space optimization as described in Rink et al (76). The exocytosis events were preceded by a docking event, which corresponds to immobilization of the endosome and a fast increase of intensity (flash) due to the physics of TIRF microscopy, with consequent total disappearance of the signal. We filtered events with a duration of flash < 4 sec.

#### Track identification and quantification

MotionTracking was used to identify and analyze vesicles near the cell surface. In the time-lapse movie, fluorescent objects (vesicles) were detected in each frame. Each object was assigned an x-y position, and its fluorescence intensity and size were calculated. Next, objects were linked across consecutive frames to generate tracks, representing the trajectories of individual vesicles over time. Each track was characterized by parameters such as speed, direction, displacement, intensity, and area. To adjust the track break thresholds, the following parameters were considered: total score, area score, intensity score, and integral intensity score. These thresholds were critical for maintaining track accuracy. After optimization of the parameter, the tracks were generated per movie. Each track was considered to represent a vesicle positioned near the plasma membrane.

#### Fixed-cell TIRF-M

Fixed-cell TIRF-M was performed by plating cells over a gelatin-coated glass-bottom dish, followed by fixation using 4% PFA, and then proceeded for immunostaining. While imaging with a TIRF microscope, the cells were kept in 1×PBS. *Z*-stack images of the fixed samples were acquired with a *z*-step of 0.02 µm. Acquired images were analyzed as maximum intensity projection (MIP) and searched for multicolor objects (as described above) using the automated image analysis program Motiontracking.

#### Super-resolution microscopy sample preparation

BT549 or MDA-MB-231 cells expressing the protein of interest were fixed in 4% PFA for 15 min at room temperature and then proceeded for indirect immunofluorescence as mentioned above. Image was captured using Zeiss LSM 900 Airyscan or Olympus IXplore spinSR microscope sequentially with laser wavelengths 488 nm and 561 nm. *Z*-stack images of the fixed samples were acquired with a *z*-step of 0.02 µm.

For Airyscan image processing, the Airyscan Joint deconvolution (jDCV) method was used, which uses the Richardson-Lucy algorithm to achieve a resolution up to 90nm.

#### Gelatin degradation assay

Gelatin degradation assay was performed as described by Sharma et.al. (6). MDA-MB-231, BT549 and MCF10DCIS cells were trypsinized and 50,000 cells were plated on Alexa Fluor 568-labeled gelatin for 3-12h. Cells were then fixed and immunostained with DAPI. Images were captured using a Zeiss LSM780 confocal laser scanning microscope with a 40×1.4 NA oil immersion objective or an Olympus FV3000 microscope. Images were processed in the Motiontracking software, where the degradation spots were identified as objects, and the degradation index was calculated using the formula

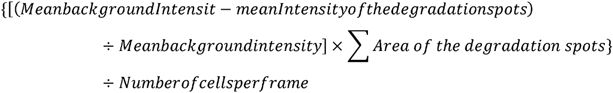

#### Generation of multicellular spheroids

Spheroids were formed as described previously in Nazari et al with minor modifications (77). Briefly, non-adherent agarose microwells were generated by pipetting 50*μ*l of pre-warmed 2% w/v agarose solution, made in PBS, into each well of a 96-well plate and allowed to solidify. MDA-MB-231 or BT549 cells were trypsinized, resuspended and 2000 cells were seeded into each well. Plates were centrifuged for 5 min at 100 ×g, 25°C to initiate aggregation of cells within each well. Plates were incubated at 37°C for 2 days to allow cells to grow and form more compact aggregates. For MDA-MB-231, on day 2, 100*μ*l of media was taken out and 100*μ*l of 10% Matrigel (v/v) (Corning, Cat#354277) was added into each well, making the final Matrigel concentration 5%. After brief centrifugation, plates were returned to the incubator and allowed to form compact spheroids for an additional 2 days.

#### Spheroid invasion into collagen and quantification

The multicellular spheroids were harvested from the microwells and added in the media. Cooled rat tail collagen type I (ThermoFischer Scientific, Cat#A1048301) was neutralized following the manufacturer’s protocol to make collagen hydrogels. The spheroids were mixed with the collagen solution for ∼ 5 spheroids/ 70*μ*l in a final collagen concentration of 2mg/mL. Next, the solution was added to a 96-well plate and allowed to gel for 30 min at 37°C. 200ul of complete media or Serum serum-free media or serum-free media containing 100ng/ml HGF was added into each well. The spheroids were allowed to invade for 48 hours, and images were captured at Day 0 and after 48 hours using a Nikon Eclipse Ti inverted microscope using the 4× objective at bright field mode. For BT549 GFP-MET WT or M1250T Mutant spheroids, doxycycline was added to complete media to make a final concentration of 50ng/ml, respectively. Spheroid images were immediately captured at Day 0 with a Nikon Eclipse Ti inverted microscope using a 4× objective at bright field mode. The spheroids were transferred to the incubator and imaged after 24 hours.

The spheroid invasion area was calculated by ImageJ. By color threshold and the wand tracing tool, the core area and the invaded area were identified, the boundary was drawn, and the area was measured. Spheroid invasion was calculated as the ratio of the invaded area to the core area and was plotted. Experiments were performed in triplicate with a minimum of 30 spheroids per condition.

#### MT1-MMP antibody uptake assay

MDA-MB-231 cells were trypsinized and counted, and 50,000 cells per well were seeded on the gelatin-coated coverslips for 24 h. After the cells had properly adhered and regained their morphology, antibody uptake was performed using anti-MT1-MMP antibody (R&D Systems, MAB9181-SP). Cells were incubated at 4°C with 10 μg ml^−1^ anti-MT1-MMP antibody diluted in serum-free L-15 medium for 1 h. The unbound antibody was removed by washing with 1× PBS. L-15 complete medium was added, and cells were shifted to 37°C to allow internalization of the surface-bound antibody for 30 minutes. Next, an acid wash (0.2 M Acetic acid and 0.5 M NaCl) was given for 4 minutes to remove the surface-bound antibody. Again, the cells were washed with 1× PBS, and complete media was added with or without HGF to allow recycling of the internalized pool. At the required time point, cells were fixed by adding 4% PFA, followed by staining with anti-EEA1 and DAPI.

The % MT1-MMP recycling was calculated using the formula:

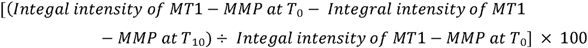

#### Western blotting

Cells were harvested and lysed in lysis buffer (50 mM Tris-HCl, pH 7.4, 150 mM NaCl, 2 mM DTT, 1% NP40, and 1 mM EDTA) supplemented with 10 μg ml^−1^ protease inhibitor cocktail (PIC, Sigma). Cell debris was removed by centrifuging at 19,000 ×g for 15 min at 4°C. Protein concentration was estimated using the Bradford assay. Samples were prepared by adding 1× SDS loading dye and boiling at 95°C. Proteins were separated by running on SDS-PAGE, then transferred to a nitrocellulose membrane (0.45 μm; GE Healthcare, Cat. 10600002). The membrane was blocked with 5% BSA, then incubated for 1 h at room temperature or overnight at 4°C with the respective primary antibody. To detect the bound antibody signal, the membrane was incubated with fluorescently labelled secondary antibody, and protein bands were detected using the Li-Cor Odyssey Infrared Scanning System.

#### Blot quantification using ImageJ

Densitometric analysis of blot images was performed using ImageJ software. Blot images were saved as 8-bit grayscale files. Using the rectangular selection tool, equal-sized boxes were drawn around each band, and the Gel Analysis function was used to generate intensity profiles. The area under each peak, corresponding to band intensity, was measured after manually defining baselines and subtracting the background signal obtained from a non-band region of the blot. Band intensities were normalized to the corresponding loading control, and relative expression levels were calculated. Quantification was performed on at least three independent experiments, and results are presented as mean ± SD.

#### MT1-MMP surface biotinylation

Surface Biotinylation was performed as described by Parveen et.al. (53). To measure the surface population, MDA-MB-231 cells were grown in a 35-mm plate. After 24h, the cells were treated with or without HGF for 10 minutes at 37°C. Then the culture medium was removed and the cells were washed with 1× PBS (6.7 mM NaHPO_4_, 3.3 mM NaH2PO_4_, and 140 mM NaCl, pH 7.2) and incubated with EZ-link Sulfo-NHS-Biotin (Invitrogen, Cat. 21217) at 4°C for 45 min. Next, to quench free biotin, cells were treated with 50 mM Tris-HCl, pH 8.0. After quenching, cells were washed with 1× PBS and lysed. The lysate was clarified and incubated with Neutravidin beads (Pierce, Cat. 29200) for 30 min at room temperature. Biotin-labelled proteins were eluted from the beads and subjected to immunoblotting. The intensity of the biotinylated MT1-MMP was quantified for HGF-treated or untreated samples using ImageJ software.

#### Co-immunoprecipitation

20µl of the protein A/G beads agarose beads were washed and incubated with 2µg of IgG or anti-MET antibody for 1h at 4°C with gentle rotation. The unbound antibodies were washed off. Subsequently the MDA-MB-231 lysate was added and further incubated for 2h at 4°C. To remove unbound proteins washes were given. The beads-antibody-protein complex containing 1× SDS sample loading buffer was boiled at 95°C. The samples were separated by SDS-PAGE and subjected to immunoblotting.

#### GFP-trap pulldown

The GBP clone was purchased from Addgene and subcloned into the pGEX-6P1 vector to express GST-tagged GBP. GFP vector or GFP fusion proteins were overexpressed in the 6-well format. After 16 h of transfection, cells were lysed in 50 mM Tris-HCl, pH 7.4, 150 mM NaCl, 2 mM DTT, 1 mM EDTA, and 1% NP-40, and cleared by centrifuging at 19,000 ×*g* at 4°C for 20 min. The cell lysates were mixed with 25 µg of purified GST–GBP bound to the glutathione Sepharose beads. The beads bound with GBP–GFP protein complexes were then washed three times with PBS, followed by elution using 1× SDS sample loading buffer. Samples were then separated on an 8% SDS-PAGE gel, followed by Western blotting.

#### His-Pulldown

His pulldown was performed as mentioned by Sharma et.al. (6). Cells expressing proteins of interest were harvested after 16 hours of transient transfection. Cells were lysed and cleared by centrifuging at 19,000 ×*g* at 4°C for 20 min. Prepared lysates were incubated with Ni-NTA agarose beads. The beads bound with His tagged protein complexes were then washed with 1× PBS and eluted. The samples were further processed for Western blotting.

#### Protein turnover experiment

The protein turnover was estimated as described by Sharma et.al. (6). 50,000 cells were seeded for 24 h before the experiment. The next day, cells were treated with or without HGF with cycloheximide at a working concentration of 10 µg ml^−1^. At different time points, cells were lysed and clarified. The cell lysates were then subjected to Western blotting.

#### PHA665752 or Pervanadate treatment

To inhibit the Tyr phosphatases, Pervanadate was added to the cells as described by Parveen et.al. (53). Briefly, 6mM of pervanadate solution was prepared by mixing 1 ml of a 20 mM sodium orthovanadate with 330 µl of 30% H_2_O_2_ and incubating for 5 min at room temperature. These cells were treated with 40 µM of pervanadate solution diluted in serum-free L-15 medium. As pervanadate/H_2_O_2_ is unstable, a fresh solution was prepared each time. Cells were treated with pervanadate for 30 min at 37°C. Cells were harvested and lysed, were analyzed by anti-phospho-MET immunoblotting (53).

#### Statistical analysis

For the statistical evaluation of the datasets from quantitative image analysis, paired or unpaired Student’s *t*-test or one-way ANOVA with a Dunnett’s multiple comparisons test was performed using GraphPad Prism 6 and 8 software version 6.01 and 8.0.2, respectively. Datasets are presented as Mean±SD or Mean± SEM. For the data that are displayed using SuperPlots, each biological replicate is distinctly color-coded and each dot represents the number of identified multicolor objects in a field of view (frame). One frame comprises 1–3 cells, contributing to a total number of cells (*n*) across the total number of frames acquired. The frame-wise values were separately pooled for each biological replicate to calculate the mean in the SuperPlots, and the SD represents the deviation of the data from these three means (78). The significance is as follows: *P*>0.05, ns (non-significant), \**P*≤0.05, \*\**P*≤0.01, \*\*\**P*≤0.001, and \*\*\*\**P*≤0.0001. The number of samples, images, and experiments used for quantification is mentioned in the respective figure legends or in figures. In all figures, *n*= number of cells and *N*= number of experiments.

## Acknowledgements

We thank Professor Philippe Chavrier [CNRS, Paris, France] for sharing the pCDNA3.1 MT1-MMP pHluorin construct and the MCF10DCIS.com line; Dr. Marc Coppolino for pEGFPN1 Mt1-MMP construct, Professor Marino Zerial (Max Planck Institute, Dresden, Germany) for sharing EEA1 antibody, pEGFP-C1 RAB4 and pEYFP C1 KIF16B construct, Prof. Subba Rao Gangi Setty (IISc, India) for GFPC1 and mCherryC1 RAB14 constructs, Professor Jim.C. Norman (University of Glasgow, UK) and Elizabeth Manning (Vanderbilt University) for pmCherry RCP, pEGFPC2 RCP; Professor Sara Courtneidge for the GFP-TKS5 construct. (Oregon Health & Science University, USA); Professor Luis S. Mayorga (IHEM-CONICET), Mendoza, Argentina, for the GFP-RAB22 construct. We acknowledge Dr. Sameena Parveen for her kind help in generating the MT1-MMP Knock-out cell line, Tanishka and Shaili for helping out with plasmid isolation. We are thankful to the Central Instrumentation Facility at IISER Bhopal, which was used to perform confocal, super-resolution and TIRF microscopy-based experiments. We also acknowledge the FIST facility at IISER Bhopal, supported by the DST, for providing the infrastructure to conduct live-cell imaging-based experiments.

## Author contribution

This work was funded by the Science and Engineering Research Board (CRG/2023/000563). Author contributions: A. Khamari: Conceptualization, data curation, investigation, methodology, visualization, analysis, manuscript writing and editing; A. Guria: investigation, methodology, visualization, analysis; K. Tak: investigation, methodology, analysis; R. Sharma: investigation, methodology, analysis; Y. Kalaidzidis: Motiontracking analysis. S. Datta: Conceptualization, data curation, analysis, funding acquisition, investigation, methodology, project administration, resources, supervision, validation, visualization, and manuscript.

---

**Figure S1: HGF-induced MET activity promotes TNBC breast cancer invasion through invadopodia-mediated ECM degradation** (A) Lysates of MDA-MB-231, BT-549 and MCF10A DCIS were separated by SDS-PAGE and analyzed by Western blot. Membranes were probed with anti-MET and anti-Vinculin antibody. (B) Bright-field images of BT-549 spheroids embedded in the collagen matrix at 0h and 48h in the presence or absence of HGF or complete media. The ratio of the invaded area to the core area was quantified and plotted. Data points from individual biological replicates are distinctly color-coded. The error bar represents Mean ±SD, One-way ANOVA with multiple comparisons. scale bar: 100µm. **** P < 0.0001, *** P < 0.001 N=3, n=30. (C) MDA-MB-231 cells were transfected with control or MET siRNA (SMART Pool or 2 individual siRNAs). After 60 hours of transfection, cells were seeded on a gelatin568-coated glass-bottom dishes for 12h. Cells were fixed and immunostained for TKS5, and cortactin. Images were captured using a confocal microscope. Degradation index along with TKS5 and Cortactin-positive structures were identified and plotted in terms of % of scrambled. The inset shows a magnified view of the region indicated by the box. The error bar represents Mean±SD, One-way ANOVA with multiple comparisons, **** P < 0.0001, *** P < 0.001, ** P < 0.01. scale bar: 10µm. Inset: 2µm. N=3, n=200. (D) BT-549 cells were seeded on Alexa568-labeled gelatin-coated coverslip for 3h, followed by incubation with or without HGF for an additional 3h. Cells were fixed, immunostained for TKS5, cortactin, and imaged with a confocal microscope. The degradation index and number of colocalized TKS5-cortactin puncta were quantified and plotted. The inset shows a magnified view of the region indicated by the box. The error bar represents Mean ±SEM, Unpaired t-test. scale bar: 10µm. Inset: 2µm, *** P < 0.001, ** P < 0.01, N=3, n=200. (E) MCF10A DCIS cells were seeded on Alexa568-labeled gelatin-coated coverslip with or without HGF for 6h. Degradation Index was quantified and plotted. The Arrowhead represents degradation spots. Data points from individual biological replicates are distinctly color-coded. The error bar represents Mean ±SD, Unpaired t-test. scale bar: 10µm. **** P < 0.0001, N=3, n=200. (F) MCF10A DCIS cells were seeded on a gelatin-coated glass-bottom dish for 3h, followed by incubation with or without HGF for 3h. Cells were fixed and immunostained for cortactin and F-actin. Images were captured using a TIRF microscope. Actin-positive cortactin puncta were quantified and plotted. The error bar represents Mean ±SEM, Unpaired t-test. scale bar: 10µm. ** P < 0.01, N=3, n=100. (G) MDA-MB-231 cells were seeded on a gelatin-coated glass-bottom dish for 3h, followed by incubation with or without HGF for 3h. Cells were fixed and immunostained for cortactin and TKS5. Images were captured using a TIRF microscope. Percentage of cells showing invadopodia formation per frame were calculated and plotted. The error bar represents Mean ±SD, Unpaired t-test. ** P < 0.01, N=3, n=90. (H) MDA-MB-231 cells were seeded on a gelatin-coated glass bottom dish for 3h, followed by incubation with or without EGF stimulation for an additional 3h. Cells were fixed and immunostained for TKS5 and cortactin. Images were captured using a TIRF microscope. TKS5 puncta positive for cortactin were considered as invadopodia. The error bar represents Mean ±SD, Unpaired t-test. scale bar: 10µm. **** P < 0.0001, N=3, n=90 (I) MDA-MB-231 cells were treated with control or SMARTPool or 2 individual MET siRNA. The efficiency of gene silencing was estimated by immunoblotting. To analyze total MT1-MMP levels in MET-depleted cells, the membrane was probed for MT1-MMP. Actin was used as a loading control. (J) MDA-MB-231 cell treated with or without HGF in the presence of vehicle or PHA665752 for 3h, lysed and separated by SDS-PAGE. The membrane was probed for pY1234-35 MET or MET and vinculin. The numbers indicates the normalized MET of p-MET intensities. To calculate p-MET intensity, the signal intensity of p-MET was normalized with MET and further with loading control. (K) MDA-MB-231 cells were transfected with SHC002 control shRNA or 5 shRNA clones against MET. After 48 hours, cells were lysed and subjected to immunoblotting with anti-MET antibody. The numbers indicate the normalized signal intensity of MET. (L) MDA-MB-231 cells were transfected with control SHC002 or MET shRNA clone #3, and after 36h of transfection, cells were seeded on Alexa568-labeled gelatin-coated coverslip for 6h. The cells were fixed and immunostained with anti-MET antibody, Phalloidin, and DAPI.

**Figure S2: HGF promotes MET recycling that drives breast cancer invasion**

(A) BT-549 cells were seeded on gelatin-coated coverslips and allowed to adhere for 3h, followed by 3 hours with HGF stimulation. Cells were fixed, immunostained for TKS5, MET, and the Nucleus. Images were captured using the confocal microscope. The inset shows a magnified view of the region indicated by the box. Scale bar: 10µm. Inset: 2µm. (B) The ratio of the integral intensity of MET to TKS5 was calculated, and the frequency distribution was plotted. n= 800 invadopodium. (C) GFP-TKS5 transfected MDA-MB-231 cells were seeded on gelatin-coated glass-bottom dishes in the presence or absence of HGF, after 6h, fixed, and immunostained for MET. The cells were imaged using a TIRF microscope. The percentage of TKS5 puncta positive for MET was quantified and plotted. scale bar: 10µm. Unpaired t-test. *** P < 0.001, N=3, n=120. (D) MDA-MB-231 cells were seeded on gelatin-coated glass coverslips for 3h, followed by incubation with or without HGF for an additional 3h. The cells were fixed and immunostained with anti-MET, anti-cortactin antibody, and Phalloidin. The cells were imaged using a confocal microscope. Numbers of Actin-cortactin puncta positive for MET were quantified and plotted. Scale bar: 10µm. Unpaired t-test. *** P < 0.001, N=3, n=200. (E) MDA-MB-231 cells seeded on glass coverslips were stimulated with or without HGF for 2h. Cells were fixed and immunostained for MET and RAB5. The percent colocalization of MET with RAB5 was calculated. scale bar: 10µm. The error bar represents Mean ±SEM. N=3, n=200. (F) MDA-MB-231 cells were treated with HGF for a period of 0-3h in the presence of cycloheximide, lysed at the indicated time points, and subjected to immunoblotting. The numbers represent the relative MET expression at the mentioned time point. The relative MET intensity was calculated for 3 replicates and plotted. (G) BT-549 cells were seeded on gelatin-coated coverslips for 6h and fixed. Cells were immunostained for EGFR, F-actin, and cortactin. Images were captured with the confocal microscope. scale bar: 10µm. (H) BT-549 cells were seeded on gelatin-coated coverslips for 6h with or without EGF stimulation and fixed. Cells were immunostained for EGFR, F-actin, and cortactin. Images were captured with the confocal microscope. The percentage of Actin-cortactin puncta that immunostained for EGFR was quantified and plotted. The inset shows a magnified view of the region indicated by the box. scale bar: 10µm. Inset: 2µm. Unpaired t-test. ** P < 0.01, N=3, n=100. (I) WT or M1250T V5-MET expressing BT-549 cells were seeded on glass coverslips cells fixed, and immunostained for V5 and p-MET. Images were acquired with a confocal microscope. Corrected total cell fluorescence (CTCF) for V5 and p-MET was calculated using ImageJ. The CTCF of p-MET was normalized with that of V5 and plotted. scale bar: 10µm. The error bar represents Mean ±SEM, Unpaired t-test. ** P < 0.01. N=3, n=10. (J) MDA-MB-231 cells expressing V5-MET WT or mutants were seeded on Alexa568-labeled gelatin-coated coverslips for 12h. Cells were fixed, then immunostained for V5, TKS5, and the Nucleus. Cells were imaged with the confocal microscope. Gelatin degradation by the V5-MET overexpressing cells was calculated. The error bar represents Mean ±SEM, Unpaired t-test. * P < 0.05 ns P > 0.05 scale bar: 10µm. N=3, n=120 (K, K’) V5-MET WT or M1250T or LL/AA mutant expressing MDA-MB-231 cells were seeded on a gelatin-coated glass-bottom dish for 6h and fixed. Cells were immunostained for TKS5, and images were captured with the TIRF microscope. Data points from individual biological replicates are distinctly color-coded. The error bar represents Mean ±SD, Unpaired t-test. Scale. ** P < 0.01, ns P > 0.05 bar: 10µm. N=3, n=60.

**Figure S3: HGF promotes RAB4 and RAB14-mediated MET delivery to the plasma membrane.**

MDA-MB-231 cells, seeded on glass coverslips, were transfected with (A) GFP-RAB4 or (B) Cherry-RAB14. After 14h of transfection, cells were incubated for 2h with or without HGF, then fixed and immunostained for MET. Images were captured using a confocal microscope. scale bar: 10µm. (C) Graph showing the percent colocalization of MET with GFP-RAB4 and Cherry-RAB14 in the presence of absence of HGF treatment. The error bar represents Mean ±SD, One-way ANOVA. ** P< 0.01. N=3, n=200. (D, E) MDA-MB-231 cells seeded on glass coverslips were transfected with (D) GFP-RAB22 or (E) GFP-RAB11. After 14h of transfection, cells were incubated for 2h with or without HGF, then fixed and immunostained for MET. Images were captured using a confocal microscope. The percent colocalization of MET with GFP-RAB22 or GFP-RAB11 in the presence or absence of HGF treatment was calculated and plotted. Scale bar: 10µm. The error bar represents Mean ±SD. Unpaired t-test. ns P > 0.05 N=3, n=200. (F) The KD efficiency of RAB4, RAB14, and RAB22 was confirmed by quantitative RT-PCR. N = 3. Values in the graph represent means ± SEM. Paired t-test, ** P < 0.01, * P < 0.05. (G) MDA-MB-231 cells were transfected with control or RAB4, RAB14, or RAB22 siRNA. After 60 hours, cells were harvested and processed for immunoblotting to analyse total MET levels. The relative MET levels were quantified. The numbers indicate the normalized MET signal intensity. (H) BT549 cells grown on coverslips were transfected with Cherry-RAB14. Cells were treated with HGF for 2h and fixed, followed by immunostaining with antibodies against MET and EEA1. Images were captured using a confocal microscope. The white arrowhead indicate the MET-RAB14-EEA1 positive puncta, yellow arrowhead indicates the MET-RAB14 endosomes devoid of EEA1. scale bar: 10µm. (I) BT-549 cells co-expressing GFP-RAB4 and Cherry-RAB14 were fixed, immunostained for MET and imaged with the super-resolution microscope. The inset shows a magnified view of the region indicated by the box. The white arrowhead represents RAB4-MET vesicles, and the yellow arrowhead represents RAB14-MET vesicles. scale bar: 10µm, Inset: 2µm. (J) MDA-MB-231 cells transfected with GFP-RAB14 and incubated with or without HGF. Cells were fixed, immunostained for MET, and then imaged using a confocal microscope. The mean Integral intensity and area of GFP-RAB14 vesicles were calculated and plotted. The error bar represents Mean ±SEM. Unpaired t-test. **** P< 0.0001, ns P>0.05, N=3, n=200. (K) MDA-MB-231 cells were transfected with control or VPS26A siRNA. After 60 hours of transfection, cells were harvested and processed for immunoblotting to analyse total MET levels. (L) BT-549 cells expressing Cherry-RAB14 were fixed, then immunostained with anti-MET and anti-GOLGIN 97 antibodies. Images were captured using the Olympus SpinSR super-resolution microscope. The inset shows a magnified view of the region indicated by the box. scale bar: 10µm, Inset: 2µm. (M) MDA-MB-231 cells seeded on glass coverslips were incubated for 2h with or without HGF. Cells were fixed and immunostained for MET and VPS35. The percent colocalization of MET with VPS35 was calculated. scale bar: 10µm. The error bar represents Mean ±SEM. Unpaired t-test. ns P> 0.05. N = 3, n = 200. (N) MDA-MB-231 cells were transfected with control or VPS26A siRNA. After 60 hours of transfection, cells were given a pulse of HGF for 5 min, and excess HGF was washed off. The cells were incubated at 37°C for 30 min and fixed. The cells were immunostained for MET and F-actin. Images were captured in a confocal microscope, and MET localized to Actin at the cell periphery was quantified. scale bar: 10µm. The error bar represents Mean ±SD. Unpaired t-test. ns P>0.05. N=3, n=200

**Figure S4: RCP-RAB14-KIF16B trafficking axis regulates MET recycling**

(A) The frequency of alteration of RCP in 27 breast cancer datasets was analyzed using the publicly available database cBioportal. The graph shows the frequency of alterations for different breast tumor cohorts. (B) BT-549 cells seeded on glass coverslips were transfected with GFP-RCP. After 14h of transfection, cells were incubated for 2h with HGF stimulation, then fixed and immunostained for MET and EEA1. Images were captured using a confocal microscope. scale bar: 10μm. (C, C’) BT-549 cells seeded on glass coverslips were transfected with GFP-RCP S435A and Cherry-RAB14. After 14h of transfection, cells were incubated for 2h with or without HGF, then fixed and immunostained for MET. Images were captured using a confocal microscope. The percent colocalization of GFP-RCP S435A with RAB14 or RAB14-MET in the presence or absence of HGF treatment was calculated and plotted. scale bar: 10µm. The error bar represents Mean ±SD. Unpaired t-test. ** P< 0.01, ns P > 0.05 N=3, n=200. (D) BT-549 cells seeded on glass coverslips were transfected with GFP-RCP. After 14h of transfection, cells were incubated for 2h with HGF stimulation, then fixed and immunostained for MET and WASH. Images were captured using a confocal microscope. scale bar: 10μm. (E) BT-549 cells depleted with RAB14 or control siRNA-treated cells were transfected with GFP-RCP for 14 hours, followed by 2 hours of HGF stimulation. Cells were fixed and immunostained for MET and EEA1. Images were captured using a confocal microscope, and the percentage colocalization of RCP with MET was quantified and plotted. The graph represents Mean ±SD. Unpaired t-test. ns P > 0.05, N=3, n=200. (F) BT-549 cells were treated with control or siRNA against KIF16B, RCP or RAB14. The efficiency of gene silencing was estimated by qRT-PCR. (G) BT-549 cells were treated with control or RCP siRNA. The efficiency of gene silencing was estimated by immunoblotting. (H) MDA-MB-231 cells were transfected with RCP or KIF16B siRNA. After 72h cells were lysed and subjected to immunoblotting with anti-MET and anti-Actin antibody. (I) BT-549 cells were treated with control or SMARTpool siRNA against RCP. 60h after transfection, cells were transfected with Cherry-RAB14. After providing 2 hours of HGF stimulation, cells were fixed and immunostained for MET and EEA1. Images were captured using a confocal microscope. The percentage colocalization of RAB14 or MET endosomes that do not EEA1 was calculated. scale bar: 10µm. The error bar represents Mean ±SD. Paired t-test, ns P > 0.05, N=3, n=200. (J) BT-549 cells depleted of RAB14 or control siRNA-treated cells were transfected with YFP-KIF16B. After 14 hours of transfection, cells were treated with HGF, followed by fixing and immunostaining for MET and EEA1. Images were captured using a confocal microscope, and the percentage colocalization of KIF16B with MET was quantified and plotted. The graph represents Mean ±SD. Unpaired t-test. * P < 0.05 N=3, n=200. (K, K’) MDA-MB-231 cells depleted of RAB14, RCP or KIF16B were seeded on Gelatin-568 for 12h. Fixed and imaged with a confocal microscope. The degradation index was calculated and plotted. scale bar: 10µm. The error bar represents Mean ±SEM. One-way ANOVA with multiple comparison, **** P < 0.0001, N=3, n=200. (L) BT-549 cells were transfected with SMARTpool or 2 individual oligoes targeting RAB14, RCP or KIF16B. After 60h of transfection, cells were seeded on Gelatin-568 for 12h. Fixed and imaged with a confocal microscope. The degradation index was calculated and plotted. scale bar: 10µm. The error bar represents Mean ±SEM. One-way ANOVA with multiple comparison, *** P < 0.001, N=3, n=200.

**Figure S5: HGF/MET facilitates ECM degradation by recycling MT1-MMP to the surface.**

(A) MDA-MB-231 cells treated with or without 100ng/ml of HGF were lysed and subjected to immunoblotting with anti-MT1-MMP antibody. Vinculin was taken as a loading control. The number indicates the normalized MT1-MMP intensity. (A’) MDA-MB-231 cells treated with 10 µg/ml of cycloheximide, either in the presence or absence of HGF, were lysed and subjected to immunoblotting with an anti-MT1-MMP antibody. Tubulin was taken as a loading control. The number indicates the normalized MT1-MMP intensity. (B, B’) Untreated or HGF-treated MDA-MB-231 cells were subjected to surface biotinylation. HGF stimulation was given for 10 minutes, followed by biotin labeling at 4°C for 45 minutes. Washes were given to remove excess biotin; subsequently, biotin was quenched, and cells were lysed. The lysate was allowed to bind with Neutravidin beads, followed by immunoblotting. The numbers indicate the Normalized signal intensity of MT1-MMP with that of WCL. In (B’), the MT1-MMP signal intensity has been normalized with CIMPR. Scale bar: The error bar represents Mean ±SD, Paired t-test, * P < 0.05. N=4. (C) MDA-MB-231 cells seeded on coverslips were incubated with MT1-MMP antibody at 4° C. After PBS washes, cells were shifted to 37° C to allow endocytosis in the presence or absence of HGF. The cells were fixed and immunostained with anti-EEA1 antibody. Images were captured using Zeiss LSM780. The integral intensity of the MT1-MMP antibody was calculated and plotted. Scale bar: 10µm. The error bar represents Mean ±SEM, ns P > 0.05, N=3, n = 200. (D) MDA-MB-231 cells were transfected with control or MET siRNA. After 60 hours of transfection, cells were seeded on a gelatin-coated glass-bottom dish and allowed to attach for 3h, followed by incubation with or without HGF for another 3h. Cells were fixed and immunostained for TKS5, MT1-MMP, and F-actin. Images were captured using a Nikon TIRF microscope. TKS5, MT1-MMP, and Actin-positive structures were identified and plotted. The inset shows a magnified view of the region indicated by the box. The error bar represents Mean ±SEM, Unpaired t-test, *** P < 0.001. scale bar: 10µm. Inset: 2µm. N=3, n=120. (E) MDA-MB-231 cells seeded on glass coverslips were incubated with or without HGF. Cells were fixed and immunostained for MT1-MMP and MET. The percentage colocalization was quantified using Motiontracking. Scale bar: 10µm. The error bar represents Mean ±SD, Unpaired t-test. **** P<0.0001. N=3, n=200. (F) Quantification of GBP pulldown assay of GFP-MT1-MMP in 3 different cell lines. The value represents the signal intensity of trapped MET normalized by pulled-down GFP or GFP-MT1-MMP per assay. GFP and GFP-MT1-MMP bars are represented by distinct colors. (G) GFP or GFP-MT1-MMP and V5/H6-MET co-expressing HeLa cell lysates were incubated with Ni-NTA beads. Washes were given, and the pull-down samples were processed for immunoblotting with anti-GFP and anti-V5 antibodies. WCL: whole-cell lysates. (H) MDA-MB-231 cell lysate was immunoprecipitated using anti-MET antibody. Immunoprecipitated proteins were separated by SDS-PAGE and analyzed by Western blot. Membranes were probed for MT1- MMP and MET. (I) BT-549 cells were treated with control or MT1-MMP siRNA. After 60h of transfection, cells were transfected with GFP-MMP2. After 14h cells were lysed and analyzed by immunoblotting. The membrane was probed with MET, MMP2, GFP, MT1-MMP and vinculin antibody. The total level of the mature form of MET with a molecular weight of 140 kDa was analyzed in control and MT1-MMP-depleted conditions. GFP-MMP2 with a molecular weight of 100kDa has been taken as appositive control and Vinculin as a loading control. The numbers indicate the normalized signal intensities. (J) BT-549 cells expressing V5-MET WT or M1250T were lysed and subjected to immunoblotting with V5, MT1-MMP, and Actin. (K) BT-549 cells co-expressing pHluorin MT1-MMP with V5-MET WT or M1250T mutant were seeded on gelatin-coated coverslips for 6h and fixed. Cells were immunostained for V5 and TKS5. Images were captured with the confocal microscope. The number of MT1-MMP puncta localizing with TKS5 was calculated and plotted. The inset shows a magnified view of the region indicated by the box. The error bar represents Mean ±SD, Unpaired t-test. scale bar: 10µm. Inset: 2µm. ** P<0.01, * P < 0.05, N=3, n=100 (L, L’) BT-549 cells expressing WT or catalytically inactive (E240A) MT1-MMP were seeded on the gelatin-coated glass-bottom dish for 6h, then fixed and immunostained for TKS5. Images were captured using a confocal microscope. The number of TKS5 puncta in MT1-MMP-expressing cells was quantified and plotted. The error bar represents Mean ±SEM, One-way ANOVA, ** P < 0.01, * P < 0.05. scale bar: 10µm. N=3, n=150. (M) SCR and MT1-MMP knock-out MDA-MB-231 cells were seeded on gelatin-coated glass-bottom dishes for 6 hours. Cells were fixed and immunostained for TKS5 and F-actin. Images were acquired using a TIRF microscope. The number of Actin-TKS5 puncta was quantified and plotted. Scale bar: 10µm. The error bar represents Mean±SD, Unpaired t-test. **** P<0.0001. N=3, n=60. (N) BT-549 cells co-expressing V5-MET WT or M1250T with GFP or GFP-MT1-MMP were lysed and incubated with Ni-NTA beads. Washes were given to remove unbound proteins. The pulled-down samples were processed for immunoblotting with antibodies against V5 and GFP.

**Figure S6: Additional blots**

(S1A) Lysates of MDA-MB-231, BT-549 and MCF10A DCIS were separated by SDS-PAGE and analyzed by Western blot. Membranes were probed with anti-MET and anti-Vinculin antibody. (S1I) MDA-MB-231 cells were treated with control or MET siRNA. The efficiency of gene silencing was estimated by immunoblotting. To analyze total MT1-MMP levels in MET-depleted cells, the membrane was probed for MT1-MMP. Actin was used as a loading control. (S1J) MDA-MB-231 cell treated with or without HGF in the presence of vehicle or PHA665752 for 3h, lysed and separated by SDS-PAGE. The membrane was probed for pY1234-35 MET or MET and vinculin. The numbers indicates the normalized MET of p-MET intensities. To calculate p-MET intensity, the signal intensity of p-MET was normalized with MET and further with loading control. (S1K) MDA-MB-231 cells were transfected with SHC002 control shRNA or 5 shRNA clones against MET. After 48 hours, cells were lysed and subjected to immunoblotting with anti-MET antibody. The numbers indicate the normalized signal intensity of MET. (S2F) MDA-MB-231 cells were treated with HGF for a period of 0-3h in the presence of cycloheximide, lysed at the indicated time points, and subjected to immunoblotting. The numbers represent the relative MET expression at the mentioned time point. The relative MET intensity was calculated for 3 replicates and plotted. (3B’) BT-549 cells expressing WT or M1250T or LL/AA V5-MET stimulated with or without HGF were lysed and incubated with Ni-NTA beads. The pulled- down samples were separated by SDS PAGE and processed for immunoblotting with anti-V5 antibody. (3B) BT-549 cells expressing WT or the mutants of V5-MET were treated with or without HGF for 2 hours, followed by cell lysis. The lysates were subjected to immunoblotting with anti-V5 and anti-vinculin antibodies. The V5 signal intensity was quantified and normalized with Vinculin. Error bar represents Mean ±SEM. Unpaired t-test. * P < 0.05. (3C) BT-549 cells expressing WT or M1250T V5-MET were lysed and incubated with Ni-NTA beads. The pulled-down samples were separated by SDS PAGE and processed for immunoblotting with anti-V5 and anti-p-MET antibody. (S3G) MDA-MB-231 cells were transfected with control or RAB4, RAB14, or RAB22 siRNA. After 60 hours, cells were harvested and processed for immunoblotting to analyse total MET levels. The relative MET levels were quantified. The numbers indicate the normalized MET signal intensity. (S4H) MDA-MB-231 cells were transfected with RCP or KIF16B siRNA. After 72h cells were lysed and subjected to immunoblotting with anti-MET and anti-Actin antibody. (5J) RCP or KIF16B were silenced using SMARTpool siRNA in MDA-MB-231 cells. After 72h of transfection of siRNA, cells were treated with HGF for 30 minutes and proceeded for surface biotinylation. The cells were incubated with biotin at 4°C for 45 minutes. Washes were given to remove excess biotin; subsequently, biotin was quenched, and cells were lysed. The lysate was allowed to bind with Neutravidin beads and the biotinylated proteins were precipitated. The samples were separated by SDS-PAGE and processed for immunoblotting. The number indicates the MET signal normalized with MET signal intensity of the respective WCL. (S5A) MDA-MB-231 cells treated with or without 100ng/ml of HGF were lysed and subjected to immunoblotting with anti-MT1-MMP antibody. Vinculin was taken as a loading control. The number indicates the normalized MT1-MMP intensity. (S5A’) MDA-MB-231 cells treated with 10 µg/ml of cycloheximide, either in the presence or absence of HGF, were lysed and subjected to immunoblotting with an anti-MT1-MMP antibody. Tubulin was taken as a loading control. The number indicates the normalized MT1-MMP intensity. (6E) GFP or GFP-MT1-MMP expressing BT-549 cells were lysed and incubated with GST agarose beads bound to 25µg of GFP binding protein (GBP). Next washes were given to remove unbound proteins and proceeded for immunoblotting with GFP and MET. WCL: whole-cell lysates. (S5H) MDA-MB-231 cell lysate was immunoprecipitated using anti-MET antibody. Immunoprecipitated proteins were separated by SDS-PAGE and analyzed by Western blot. Membranes were probed for MT1-MMP and MET. (S5I) BT-549 cells were treated with control or MT1-MMP siRNA. After 60h of transfection, cells were transfected with GFP-MMP2. After 14h cells were lysed and analyzed by immunoblotting. The membrane was probed with MET, MMP2, GFP, MT1-MMP and vinculin antibody. The total level of the mature form of MET with a molecular weight of 140 kDa was analyzed in control and MT1-MMP-depleted conditions. GFP-MMP2 with a molecular weight of 100kDa has been taken as appositive control and Vinculin as a loading control. The numbers indicate the normalized signal intensities. (S5J) BT-549 cells expressing V5-MET WT or M1250T were lysed and subjected to immunoblotting with V5, MT1-MMP, and Actin. (S5N) BT-549 cells co-expressing V5-MET WT or M1250T with GFP or GFP-MT1-MMP were lysed and incubated with Ni-NTA beads. Washes were given to remove unbound proteins. The pulled-down samples were processed for immunoblotting with antibodies against V5 and GFP.

**MOVIE 1. HGF promotes spheroid invasion**

Spheroids were formed using MDA-MB-231 cells and embedded in collagen for 24h. Bright field images of the Spheroids invading into the collagen matrix in the presence or absence of HGF stimulation (Fig. 1A). 1 frame/15mins for 24 hours. Scale bar: 100µm.

**MOVIE 2. HGF promotes the surface delivery of MET**

BT-549 cells transfected with GFP-MET were seeded on the gelatin-coated glass-bottom dish and imaged using a TIRF microscope for 30s without intervals (Fig. 2H). Scale bar: 10µm, Exposure time (488): 900ms.

**MOVIE 3. RAB4 and RAB14 facilitate MET trafficking to the surface**

RAB4, RAB14, or RAB22 and control siRNA-treated BT-549 cells were transfected with GFP-MET. The cells were seeded on the gelatin-coated glass-bottom dish and imaged using a TIRF microscope for 30s without intervals. (Fig. 3D-E). Scale bar: 10µm, Exposure time (488): 900ms,

**MOVIE 4. GFP-RAB14 endosomes make tubular protrusions**

BT-549 cells expressing GFP-RAB14 were seeded on a glass-bottom dish and imaged using a confocal microscope (Fig. 5C). Scale bar: 10µm, Inset: 2µm. Exposure time: 80ms, 1frame/ 0.4s for 1 min

**MOVIE 5. RCP and RAB14 localize to the endosomal tubules**

BT-549 cells co-expressing GFP-RCP and Cherry-RAB14 were seeded on a glass-bottom dish and imaged using a confocal microscope (Fig. 5D). Scale bar: 10µm, Exposure time: 80ms, 1frame/ 0.4s for 1 min

**MOVIE 6. KIF16B promotes RAB14 endosomal tubulation**

BT-549 cells co-expressing Cherry-RAB14 with YFP-KIF16B WT or YFP-KIF16B S109A were seeded on a glass-bottom dish and imaged using a confocal microscope (Fig. 5I). Scale bar: 10µm, Exposure time: 80ms, 1frame/ 2.25s for 90sec.

**MOVIE 7 (A, B). HGF promotes MT1-MMP surface delivery**

MDA-MB-231 cells expressing pH-MT1-MMP were seeded on the gelatin-coated glass-bottom dishes in the presence (7B) or absence of HGF (7A). Time-lapse images were captured using a TIRF microscope without an interval for 300 frames. Scale bar: 10µm, Exposure time: 500ms,

**MOVIE 8. MET and MT1-MMP co-traffic at the cell surface**

MDA-MB-231 cells co-expressing GFP-MET and Cherry-MT1-MMP were seeded on the gelatin-coated glass-bottom dishes. Time-lapse imaging was done using a TIRF microscope (Fig. 6F). Scale bar:10µm, Exposure time: 900ms (TIRF 488), 500 (TIRF 568), 1 frame/3s for 30s.

