## Supplementary material for "RAB14-dependent tubulovesicular recycling directs MET to invadopodia, promoting TNBC cell invasion": S1

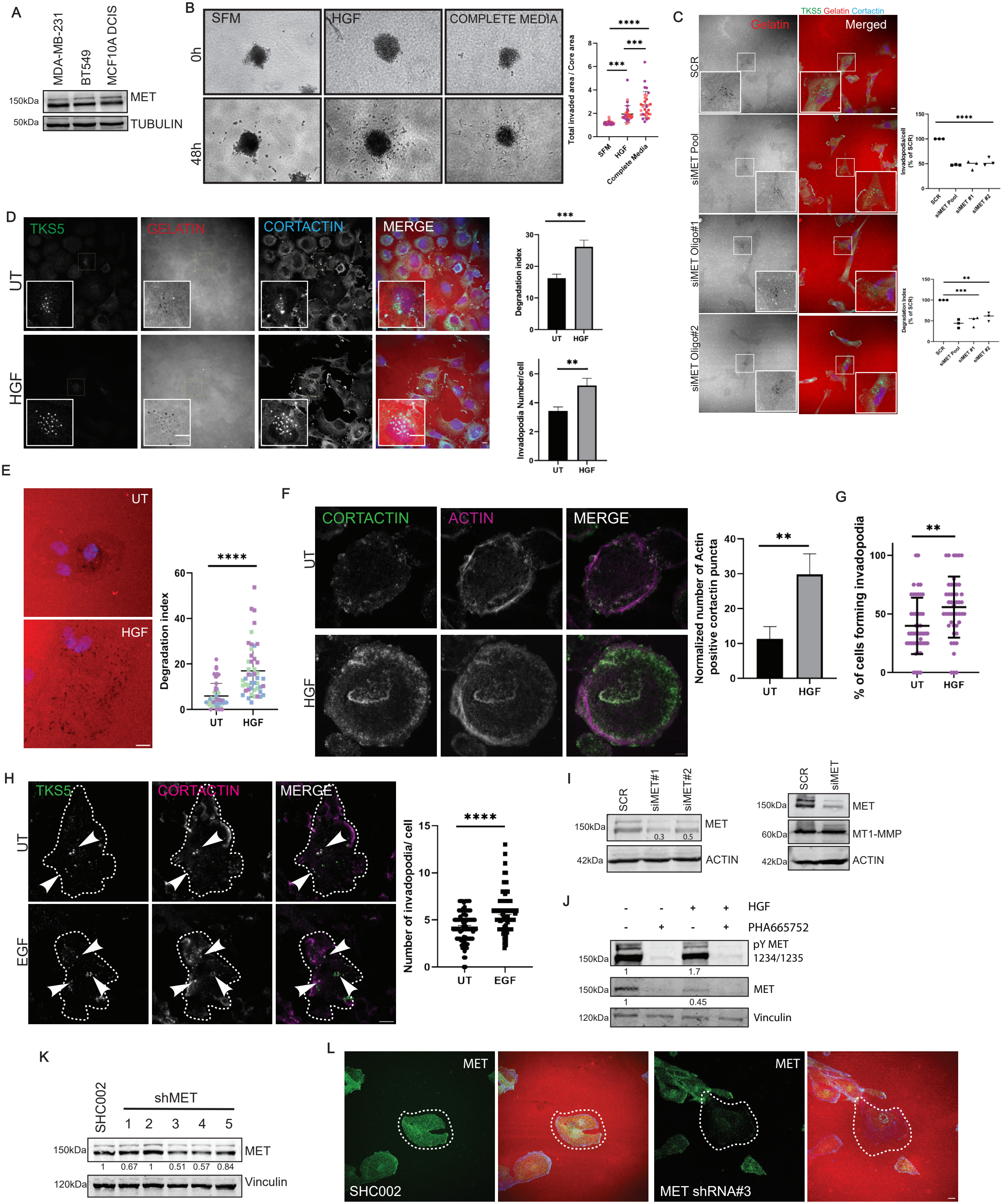
A
MDA-MB-231
BT549
MCF10A DCIS
150kDa
MET
50kDa
TUBULIN
B
SFM
HGF
COMPLETE MEDIA
0h
48h
Total invaded area / Core area
SFM
HGF
Complete Media
C
TKS5 Gelatin Cortactin
Gelatin
Merged
SCR
siMET Pool
siMET Oligo#1
siMET Oligo#2
Invadopodia/cell (% of SCR)
Degradation index (% of SCR)
D
TKS5
GELATIN
CORTACTIN
MERGE
UT
HGF
Degradation index
Invadopodia Number/cell
E
UT
HGF
Degradation index
F
CORTACTIN
ACTIN
MERGE
Normalized number of Actin positive cortactin puncta
G
% of cells forming invadopodia
H
TKS5
CORTACTIN
MERGE
EGF
Number of invadopodia/ cell
I
SCR
siMET#1
siMET#2
150kDa
MET
0.3
0.5
42kDa
ACTIN
siMET
60kDa
MT1-MMP
J
HGF
PHA665752
pY MET 1234/1235
1
1.7
0.45
120kDa
Vinculin
K
SHC002
shMET
1 2 3 4 5
1 0.67 1 0.51 0.57 0.84
L
MET
SHC002
MET shRNA#3
