## Supplementary material for "RAB14-dependent tubulovesicular recycling directs MET to invadopodia, promoting TNBC cell invasion": S2

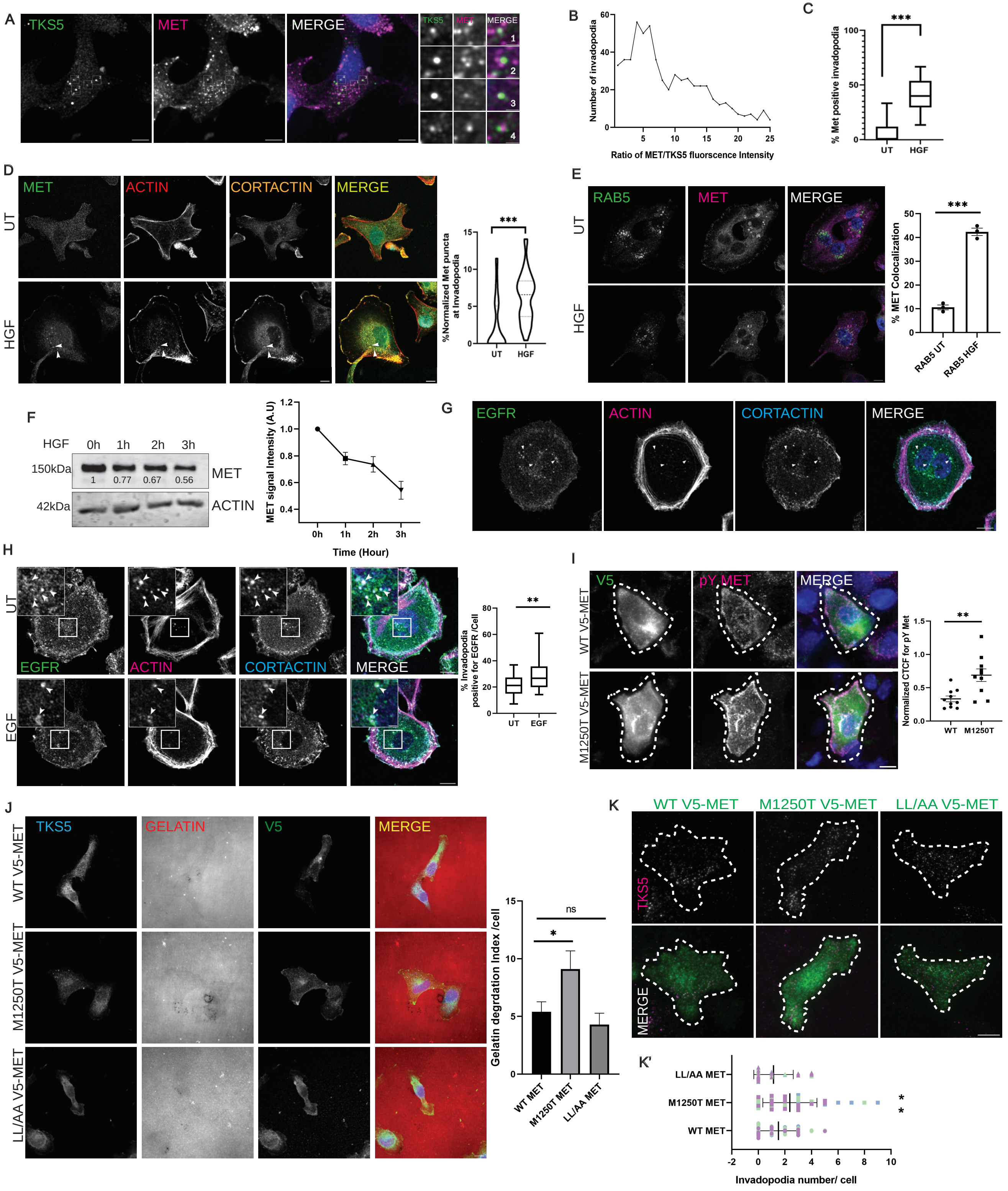

A
TKS5
MET
MERGE
B
Number of invadopodia
Ratio of MET/TKS5 fluorscence Intensity
C
% Met positive invadopodia
UT
HGF
D
MET
ACTIN
CORTACTIN
MERGE
%Normalized Met puncta at Invadopodia
E
RAB5
MET
MERGE
% MET Colocalization
RAB5 UT
RAB5 HGF
F
HGF
0h
1h
2h
3h
150kDa
MET
1
0.77
0.67
0.56
42kDa
ACTIN
MET signal Intensity (A.U)
Time (Hour)
G
EGFR
ACTIN
CORTACTIN
MERGE
H
EGF
% Invadopodia positive for EGFR /Cell
I
V5
pY MET
WT V5-MET
M1250T V5-MET
Normalized CTCF for pY Met
WT
M1250T
J
TKS5
GELATIN
V5
LL/AA V5-MET
Gelatin degrdation Index /cell
WT MET
M1250T MET
LL/AA MET
ns
K
K'
Invadopodia number/ cell
