## Supplementary material for "RAB14-dependent tubulovesicular recycling directs MET to invadopodia, promoting TNBC cell invasion": S3

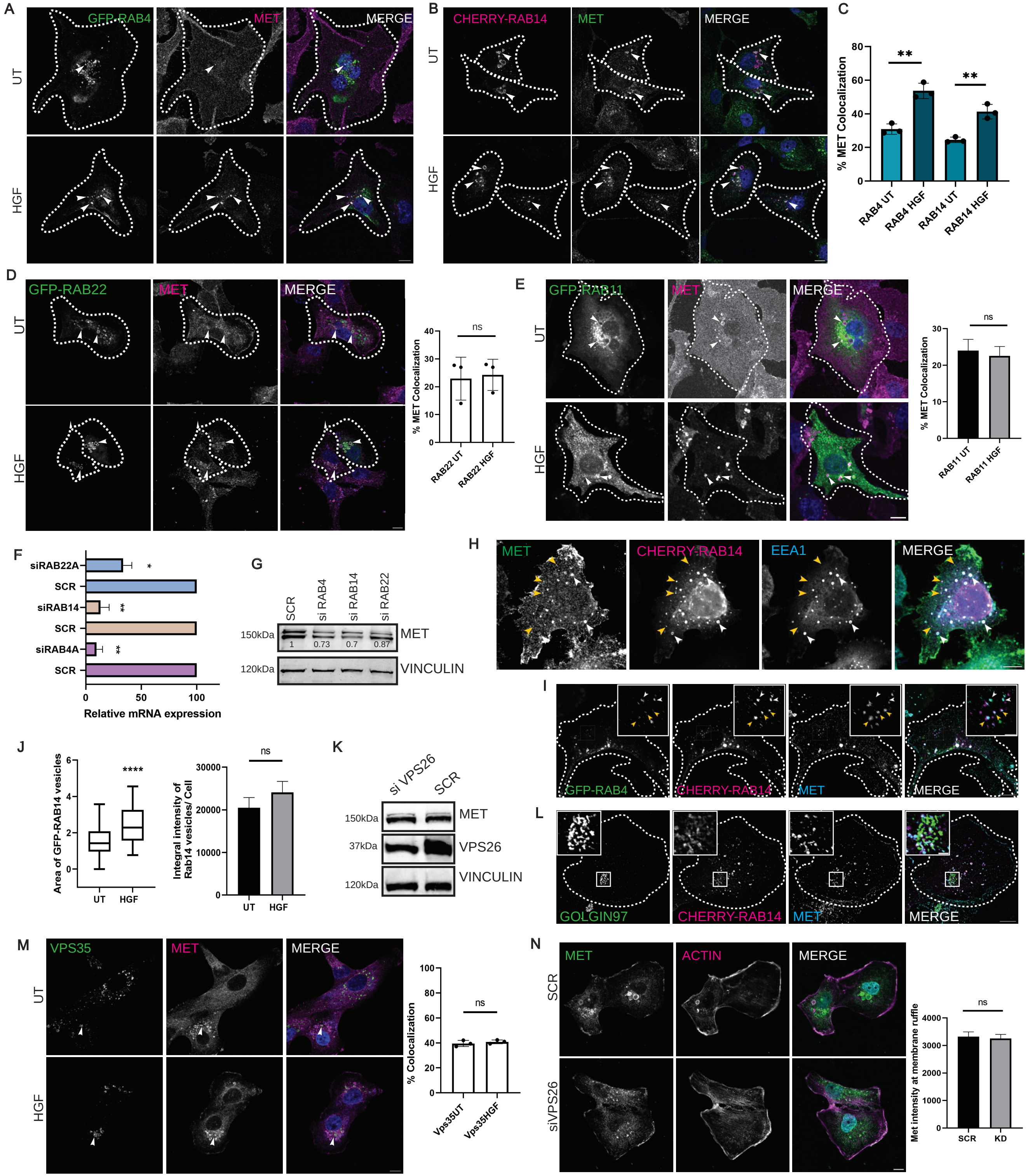
A
GFP-RAB4
MET
MERGE
UT
HGF
B
CHERRY-RAB14
MET
MERGE
UT
HGF
C
% MET Colocalization
**
**
RAB4 UT
RAB4 HGF
RAB14 UT
RAB14 HGF
D
GFP-RAB22
MET
MERGE
UT
HGF
% MET Colocalization
ns
RAB22 UT
RAB22 HGF
E
GFP-RAB11
MET
MERGE
UT
HGF
% MET Colocalization
ns
RAB11 UT
RAB11 HGF
F
siRAB22A
SCR
siRAB14
SCR
siRAB4A
SCR
Relative mRNA expression
G
SCR
si RAB4
si RAB14
si RAB22
150kDa
MET
1
0.73
0.7
0.87
120kDa
VINCULIN
H
MET
CHERRY-RAB14
EEA1
MERGE
I
GFP-RAB4
CHERRY-RAB14
MET
MERGE
J
Area of GFP-RAB14 vesicles
****
UT
HGF
Integral intensity of Rab14 vesicles/ Cell
ns
UT
HGF
K
si VPS26
SCR
150kDa
MET
37kDa
VPS26
120kDa
VINCULIN
L
GOLGIN97
CHERRY-RAB14
MET
MERGE
M
VPS35
MET
MERGE
UT
HGF
% Colocalization
ns
Vps35UT
Vps35HGF
N
MET
ACTIN
MERGE
SCR
siVPS26
Met intensity at membrane ruffle
ns
SCR
KD
