## Supplementary material for "RAB14-dependent tubulovesicular recycling directs MET to invadopodia, promoting TNBC cell invasion": S4

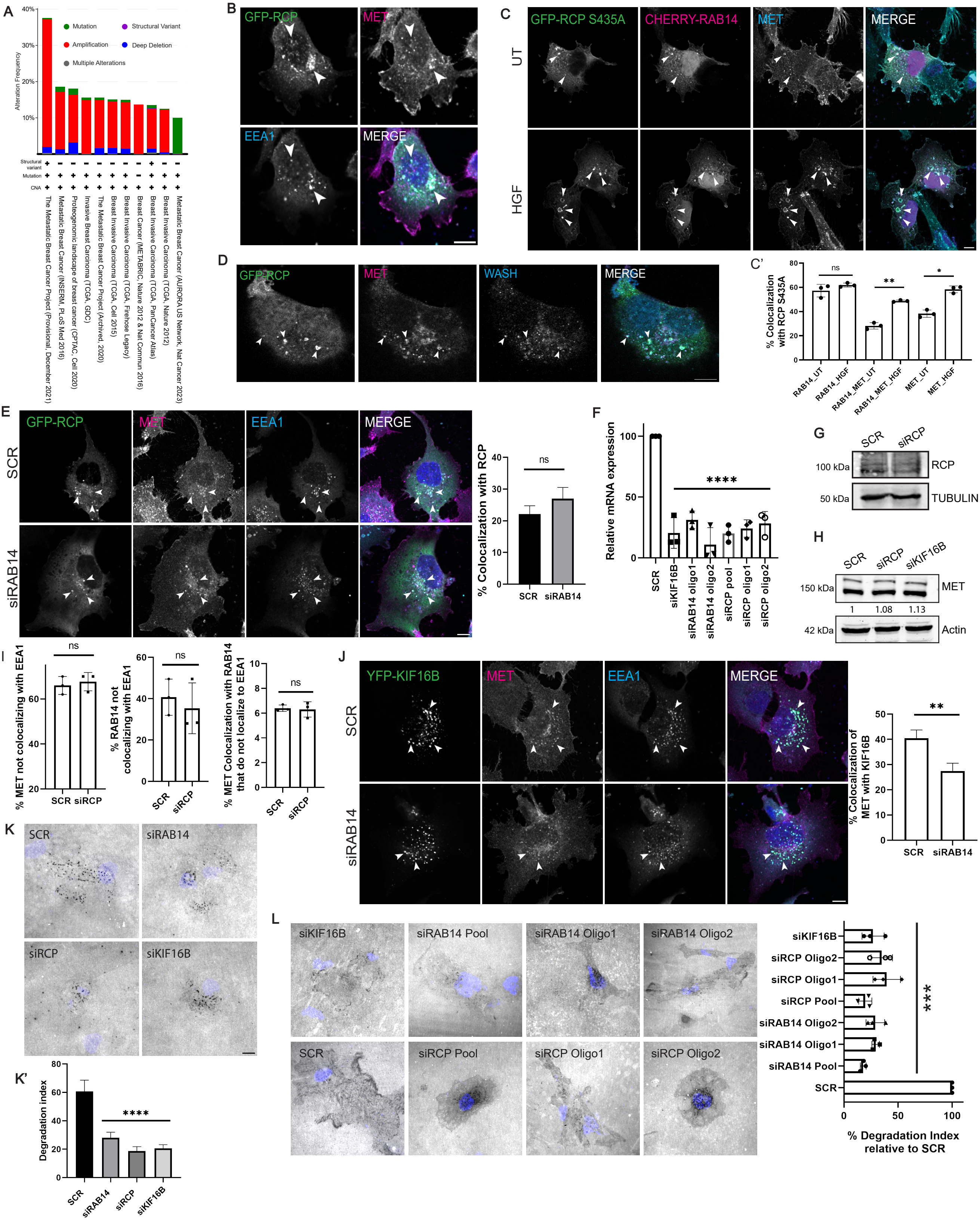

A
Mutation
Structural Variant
Amplification
Deep Deletion
Multiple Alterations
Alteration Frequency
40%
30%
20%
10%
Structural variant
Mutation
CNA
The Metastatic Breast Cancer Project (Provisional, December 2021)
Metastatic Breast Cancer (INSERM, PLoS Med 2016)
Proteogenomic landscape of breast cancer (CPTAC, Cell 2020)
Invasive Breast Carcinoma (TCGA, GDC)
The Metastatic Breast Cancer Project (Archived, 2020)
Breast Invasive Carcinoma (TCGA, Cell 2015)
Breast Invasive Carcinoma (TCGA, Firehose Legacy)
Breast Cancer (METABRIC, Nature 2012 & Nat Commun 2016)
Breast Invasive Carcinoma (TCGA, PanCancer Atlas)
Breast Invasive Carcinoma (TCGA, Nature 2012)
Metastatic Breast Cancer (AURORA US Network, Nat Cancer 2023)
B
GFP-RCP
MET
EEA1
MERGE
C
GFP-RCP S435A
CHERRY-RAB14
MET
MERGE
UT
HGF
C'
% Colocalization with RCP S435A
ns
**
*
RAB14_UT
RAB14_HGF
RAB14_MET_UT
RAB14_MET_HGF
MET_UT
MET_HGF
D
GFP-RCP
MET
WASH
MERGE
E
GFP-RCP
MET
EEA1
MERGE
SCR
siRAB14
% Colocalization with RCP
ns
SCR
siRAB14
F
Relative mRNA expression
****
SCR
siKIF16B
siRAB14 oligo1
siRAB14 oligo2
siRCP pool
siRCP oligo1
siRCP oligo2
G
SCR
siRCP
100 kDa
RCP
50 kDa
TUBULIN
H
SCR
siRCP
siKIF16B
150 kDa
MET
1
1.08
1.13
42 kDa
Actin
I
% MET not colocalizing with EEA1
ns
SCR
siRCP
% RAB14 not colocalizing with EEA1
ns
% MET Colocalization with RAB14 that do not localize to EEA1
ns
J
YFP-KIF16B
MET
EEA1
MERGE
SCR
siRAB14
% Colocalization of MET with KIF16B
**
SCR
siRAB14
K
SCR
siRAB14
siRCP
siKIF16B
K'
Degradation index
****
SCR
siRAB14
siRCP
siKIF16B
L
siKIF16B
siRAB14 Pool
siRAB14 Oligo1
siRAB14 Oligo2
SCR
siRCP Pool
siRCP Oligo1
siRCP Oligo2
***
% Degradation Index relative to SCR
