## Supplementary material for "RAB14-dependent tubulovesicular recycling directs MET to invadopodia, promoting TNBC cell invasion": S5

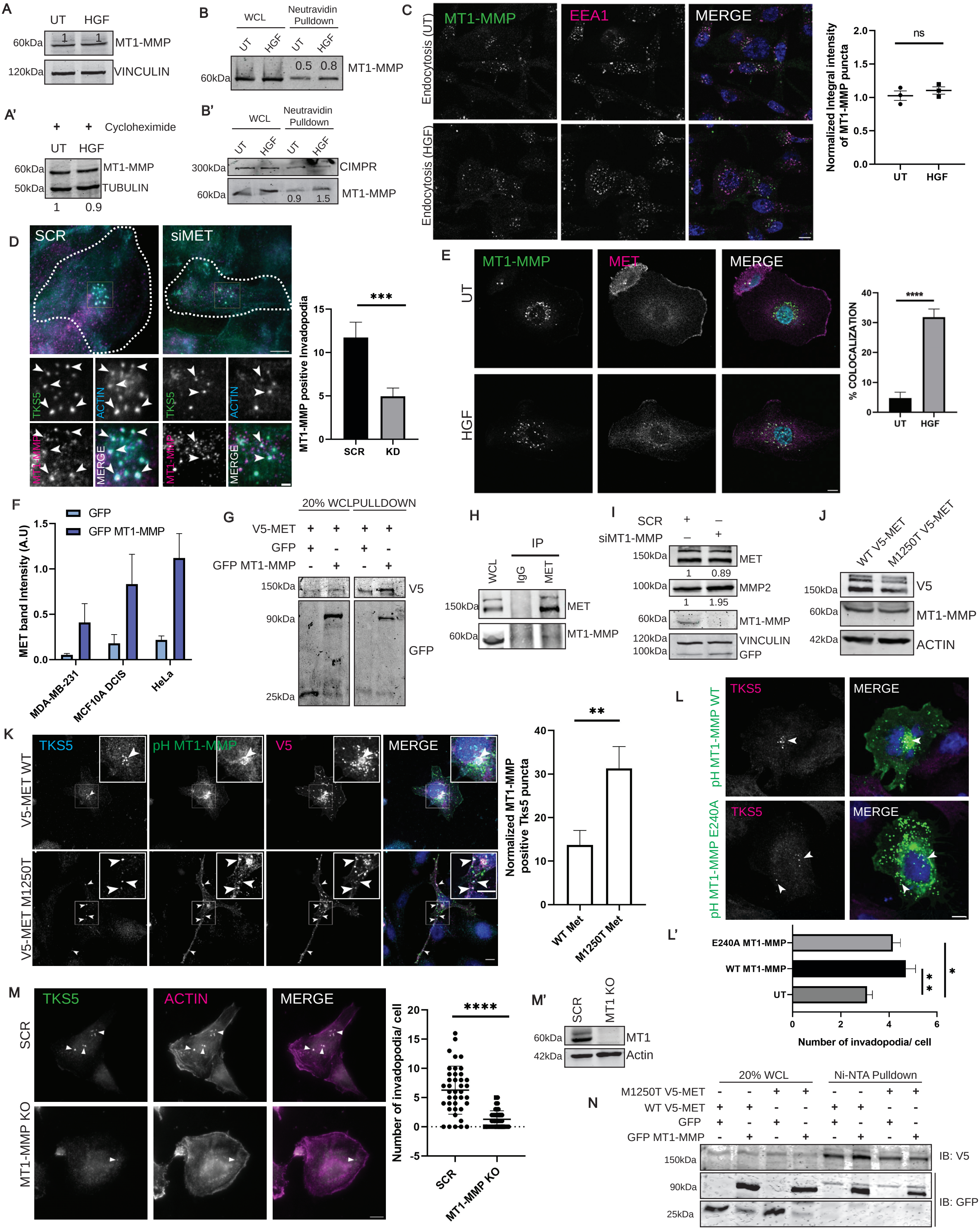
A
UT
HGF
60kDa
1
1
MT1-MMP
120kDa
VINCULIN
A'
+ Cycloheximide
UT
HGF
60kDa
MT1-MMP
50kDa
TUBULIN
1
0.9
B
WCL
Neutravidin Pulldown
UT
HGF
60kDa
0.5
0.8
MT1-MMP
B'
WCL
Neutravidin Pulldown
300kDa
CIMPR
60kDa
0.9
1.5
MT1-MMP
C
MT1-MMP
EEA1
MERGE
Endocytosis (UT)
Endocytosis (HGF)
ns
Normalized Integral intensity of MT1-MMP puncta
2.0
1.5
1.0
0.5
0.0
UT
HGF
D
SCR
siMET
TKS5
ACTIN
MT1-MMP
MERGE
***
MT1-MMP positive invadopodia
15
10
5
0
SCR
KD
E
MT1-MMP
MET
MERGE
UT
HGF
****
% COLOCALIZATION
40
30
20
10
0
UT
HGF
F
GFP
GFP MT1-MMP
MET band Intensity (A.U)
1.5
1.0
0.5
0.0
MDA-MB-231
MCF10A DCIS
HeLa
G
20% WCL
PULLDOWN
V5-MET
GFP
GFP MT1-MMP
150kDa
V5
90kDa
GFP
25kDa
H
IP
WCL
IgG
MET
150kDa
MET
60kDa
MT1-MMP
I
SCR
siMT1-MMP
150kDa
MET
1
0.89
100kDa
MMP2
1
1.95
60kDa
MT1-MMP
120kDa
VINCULIN
100kDa
GFP
J
WT V5-MET
M1250T V5-MET
150kDa
V5
60kDa
MT1-MMP
42kDa
ACTIN
K
TKS5
pH MT1-MMP
V5
MERGE
V5-MET WT
V5-MET M1250T
**
Normalized MT1-MMP positive Tks5 puncta
40
30
20
10
0
WT Met
M1250T Met
L
TKS5
MERGE
pH MT1-MMP WT
pH MT1-MMP E240A
L'
E240A MT1-MMP
WT MT1-MMP
UT
*
*
*
0
2
4
6
Number of invadopodia/ cell
M
TKS5
ACTIN
MERGE
SCR
MT1-MMP KO
****
Number of invadopodia/ cell
20
15
10
5
0
-5
SCR
MT1-MMP KO
M'
SCR
MT1 KO
60kDa
MT1
42kDa
Actin
N
20% WCL
Ni-NTA Pulldown
M1250T V5-MET
WT V5-MET
GFP
GFP MT1-MMP
150kDa
IB: V5
90kDa
IB: GFP
25kDa
