## Supplementary material for "RAB14-dependent tubulovesicular recycling directs MET to invadopodia, promoting TNBC cell invasion": S6

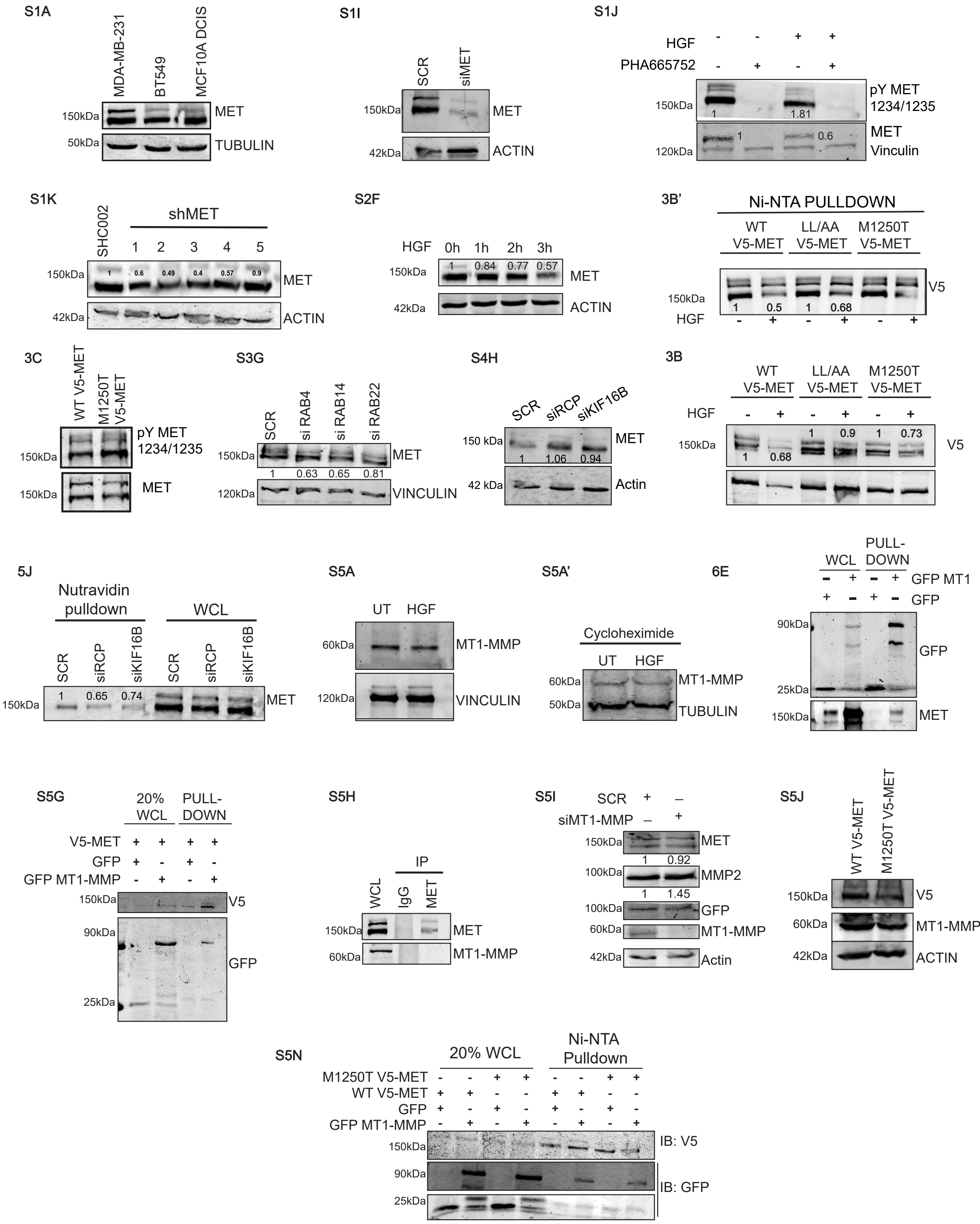
S1A
MDA-MB-231
BT549
MCF10A DCIS
150kDa
MET
50kDa
TUBULIN
S1I
SCR
siMET
150kDa
MET
42kDa
ACTIN
S1J
HGF - - + +
PHA665752 - + - +
150kDa
pY MET 1234/1235
1
1.81
MET
Vinculin
1
0.6
120kDa
S1K
SHC002
shMET
1 2 3 4 5
150kDa
1 0.6 0.49 0.4 0.57 0.9
MET
42kDa
ACTIN
S2F
HGF 0h 1h 2h 3h
150kDa
1 0.84 0.77 0.57
MET
42kDa
ACTIN
3B'
Ni-NTA PULLDOWN
WT V5-MET
LL/AA V5-MET
M1250T V5-MET
V5
150kDa
1 0.5 1 0.68
HGF - + - + - +
3C
WT V5-MET
M1250T V5-MET
pY MET 1234/1235
150kDa
MET
150kDa
S3G
SCR
si RAB4
si RAB14
si RAB22
150kDa
MET
1 0.63 0.65 0.81
120kDa
VINCULIN
S4H
SCR
siRCP
siKIF16B
150 kDa
MET
1 1.06 0.94
42 kDa
Actin
3B
WT V5-MET
LL/AA V5-MET
M1250T V5-MET
HGF - + - + - +
150kDa
1 0.68
1 0.9 1 0.73
V5
5J
Nutravidin pulldown
WCL
SCR
siRCP
siKIF16B
SCR
siRCP
siKIF16B
150kDa
1 0.65 0.74
MET
S5A
UT HGF
60kDa
MT1-MMP
120kDa
VINCULIN
S5A'
Cycloheximide
UT HGF
60kDa
MT1-MMP
50kDa
TUBULIN
6E
WCL
PULL-DOWN
GFP MT1
GFP
90kDa
GFP
25kDa
150kDa
MET
S5G
20% WCL
PULL-DOWN
V5-MET + + + +
GFP + - + -
GFP MT1-MMP - + - +
150kDa
V5
90kDa
GFP
25kDa
S5H
IP
WCL
IgG
MET
150kDa
MET
60kDa
MT1-MMP
S5I
SCR + –
siMT1-MMP – +
150kDa
MET
1 0.92
100kDa
MMP2
1 1.45
100kDa
GFP
60kDa
MT1-MMP
42kDa
Actin
S5J
WT V5-MET
M1250T V5-MET
150kDa
V5
60kDa
MT1-MMP
42kDa
ACTIN
S5N
20% WCL
Ni-NTA Pulldown
M1250T V5-MET - - + + - - + +
WT V5-MET + + - - + + - -
GFP + - + - + - + -
GFP MT1-MMP - + - + - + - +
150kDa
IB: V5
90kDa
IB: GFP
25kDa
