## Supplementary material for "RAB14-dependent tubulovesicular recycling directs MET to invadopodia, promoting TNBC cell invasion": TABLE

### TABLE 1: LIST OF PRIMERS

| NAME | SEQUENCE |
|---|---|
| **CLONING PRIMERS** | |
| **MET Fwd** | CCGCTCGAGATGAAGGCCCCCGCTGTGCTTGCACCTGGCATCCTCGTG |
| **MET Rev** | CCCAAGCTTTGATGTCTCCCAGAAGGAGGCTGG |
| **MMP14 E240A Fwd** | GCTGTGCACGCGCTGGGCCATGC |
| **MMP14 E240A Rev** | ATGGCCCAGCGCGTGCACAGCCACC |
| **LL/AA fwd** | GTATCCTGCTGCGTTGTCATCAGAAGATAACGCTGATGATGAGGTGGACAC |
| **LL/AA Rev** | GATGACAACGCAGCAGGATACGGAGCGACACATTTTACGTTCACATAAGTAGCGTT |
| **M1250T Fwd** | GTGAAGTGGACGGCTTTGGAAAGTCTGCAAACTC |
| **M1250T Rev** | CAGACTTTCCAAAGCCGTCCACTTCACTGG |
| **RT-PCR PRIMERS** | |
| **RAB4 FWD** | CACCAGCCGAGAAACCTACA |
| **RAB4 REV** | GGGTCCAGCTCACCTGATTC |
| **RAB14 FWD** | GCGAGTGCAAAAACGGGAG |
| **RAB14 REV** | CAGCCTTCTCTCTGGGGTTG |
| **RAB22 FWD** | TGGCCTCTCCCTTCTCAACT |
| **RAB22 REV** | TACACCTGTATCCCCGAGCA |
| **RCP FWD** | GCTGTCAAGCCCCGACTTC |
| **RCP REV** | TATGCAAATGCAGGGTCCGA |
| **KIF16B FWD** | CTCTTTGGGCGCAAATCCAG |
| **KIF16B REV** | ATTCAAGGGCAGCAAGCTCT |

### TABLE 2: siRNA SEQUENCE

| Gene Name | siRNA sequence |
|---|---|
| **MET** | GAACUGGUGUCCCGGAUAU |
| **MET** | GAACAGCGAGCUAAAUAUA |
| **MET** | GAGCCAGCCUGAAUGAUGA |
| **MET** | GUAAGUGCCCGAAGUGUAA |
| **RAB4A** | GCUCAGGAGUGUGGUUGUU |
| **RAB4A** | UACAAUGCGCUUACUAAUU |
| **RAB4A** | GAUAAUAAAUGUUGGUGGU |
| **RAB4A** | GAACGAUUCAGGUCCGUGA |
| **VPS26A** | GCUAGAACACCAAGGAAUU |
| **VPS26A** | UAAAGUGACAAUAGUGAGA |
| **VPS26A** | UGAGAUCGAUAUUGUUCUU |
| **VPS26A** | CCACCUAUCCUGAUGUUAA |
| **Scrambled** | UGGUUUACAUGUCGACUAA |
| **Scrambled** | UGGUUUACAUGUUGUGUGA |
| **Scrambled** | UGGUUUACAUGUUUUCUGA |
| **Scrambled** | UGGUUUACAUGUUUUCCUA |
| **RAB14** | GCUCUUAUGGUCUAUGAUA |
| **RAB14** | CAACUGCACCAUACAACUA |
| **RAB14** | GAAAAUGGCUUAUUGUUCC |
| **RAB14** | GUACAAGAAUAAUCGAAGU |
| **MT1-MMP** | GGAUGGACACGGAGAAUUU |
| **MT1-MMP** | GGAAACAAGUACUACCGUU |
| **MT1-MMP** | GGUCUCAAAUGGCAACAUA |
| **MT1-MMP** | GAUCAAGGCCAAUGUUCGA |
| **RCP** | AAACAGAAGGAAACGAUAA |
| **RCP** | GGAAAGAUGUAAAUCAGCA |
| **RCP** | AGUGAGAACUUGAACAAUG |

| | |
|---|---|
| **RCP** | CCACCAAGGUUGCUAACUG |
| **RAB22** | GGACUACGCCGACUCUAUU |
| **RAB22** | GAAGAAUUCCAUCCACUGA |
| **RAB22** | GAAACAACCUCUGCGAAUU |
| **RAB22** | GCAGUUUGAUUAUCCGAUU |
| **KIF16B** | GAUGAGACAUGGACUGUAU |
| **KIF16B** | GAGUACAGGCUGCAAUAUA |
| **KIF16B** | CAAAGACGCCUUCAGGAUU |
| **KIF16B** | CGAAACAUACCAUUUGUGA |

### TABLE 3: shRNA SEQUENCE

| Gene Name | shRNA sequence |
|---|---|
| **MET** | CCGGTCGATATTCTTTGCTCCTTGCCTCGAGGCAAGGAGCAAAGAATATCGATTTTT |
| **MET** | CCGGCCAATTTATCAGGAGGTGTTTCTCGAGAAACACCTCCTGATAAATTGGTTTTT |
| **MET** | CCGGCAGAAGTGATTGTGGAGCATACTCGAGTATGCTCCACAATCACTTCTGTTTTT |
| **MET** | CCGGCTGTTATTACTACTTGGGTTTCTCGAGAAACCCAAGTAGTAATAACAGTTTTT |
| **MET** | CCGGCACTGCTTTAATAGGACACTTCTCGAGAAGTGTCCTATTAAAGCAGTGTTTTT |
| **SHC002** | CCGGCAACAAGATGAAGAGCA CCAACTCGAGTTGGTGCTCTTC ATCTTGTTGTTTTT |
